# Morphometrics of complex cell shapes: Lobe Contribution Elliptic Fourier Analysis (LOCO-EFA)

**DOI:** 10.1101/157842

**Authors:** Yara E. Sánchez-Corrales, Matthew Hartley, Jop van Rooij, Athanasius F. M. Marée, Verônica A. Grieneisen

## Abstract

Quantifying cell morphology is fundamental to the statistical study of cell populations, and can help us unravel mechanisms underlying cell and tissue morphogenesis. Current methods, however, require extensive human intervention, are highly sensitive to parameter choice, or produce metrics that are difficult to interpret biologically. We therefore developed a novel method, Lobe Contribution Elliptical Fourier Analysis (LOCO-EFA), which generates from digitalised cell outlines meaningful descriptors that can be directly matched to morphological features. We show that LOCO-EFA provides a tool to phenotype efficiently and objectively populations of cells by applying it to the complex shaped pavement cells of *Arabidopsis thaliana* wild type and *speechless* leaves. To further validate our method, we analysed computer-generated tissues, where cell shape can be specified in a controlled manner. LOCO-EFA quantifies deviations between the specified shape that an individual *in silico* cell takes up when in isolation and the resultant shape when they are allowed to interact within a confluent tissue, thereby assessing the role of cell-cell interactions on population cell shape distributions.

**Summary statement**

Novel method (LOCO-EFA) quantifies complex cell shapes, extracting meaningful biological features such as protrusion number and amplitude; here shown for plant pavement cells and validated on *in silico* tissues.

## Introduction

Cell geometry has long fascinated biologists (Thompson, 1917). This interest is driven by a wide range of underlying scientific questions. For instance, cell shape changes can be linked to physiological responses of cells, such as membrane protrusions during apoptosis and migration (Charras and Paluch, 2008); they are involved in controlling tissue homeostasis (Marinari et al., 2012; Veeman and Smith, 2013) and underlie cell behaviour such as chemotaxis to ensure survival (Keren et al., 2008; Driscoll et al., 2012). The cell shape itself also influences intracellular processes such as microtubule organisation (Gomez et al., 2016) and the stress patterns in plant epithelia (Sampathkumar et al., 2014); it indirectly positions the plane of cell division (Minc et al., 2011) and can even determine the way a flower attracts pollinators (Noda et al., 1994). Given the rich diversity of processes in which cell shape plays a decisive role, either actively or passively, it is not surprising that the qualitative and quantitative study of cell shape characteristics, cell morphometrics, has become important in developmental biology, enriching molecular and genetic techniques. In parallel, recent advances in imaging technology, including microscopy, usage of fluorescent proteins and imaging software, allow us to collect and access remarkable amounts of cell morphological data, which in turn urgently calls for analytic tools for the extraction of meaningful cell shape information (Zhong et al., 2012). In stark contrast to the technological advances on the imaging front, there are relatively few automatic and quantitative tools available to analyse complex cell shapes (Rajaram et al., 2012; Ljosa et al., 2012; Ivakov and Persson, 2013). This gap reflects the non-trivial nature of this task: cell shape is often irregular and variable, making it very difficult to establish universal criteria encompassing cell geometry.

To illustrate the issues involved in capturing quantitative aspects of complex cell shapes, consider the case of pavement cells in the plant epidermis (Figure 1A, B) or amnioserosa cells in the Drosophila embryo (Figure 1C). Pavement cells present a striking development, requiring multiple locally divergent growth fronts within each cell, which are coordinated amongst neighbouring cells; amnioserosa cells present dynamical cell shape fluctuations within a confluent tissue. The resultant jigsaw-puzzle-like features of such cells illustrate the challenges in quantitatively characterising cell shape: 1) their complex non-holomorphic geometry cannot be captured in a meaningful way with traditional shape metrics (see below); and 2) the lack of recognisable landmarks (neither these plant nor animal cells have distinct features to serve as reference points), renders a myriad of shape statistics that have been developed in other fields (such as principal component and Procrustes analysis) effectively useless (Klingenberg, 2010).

**Figure 1:**
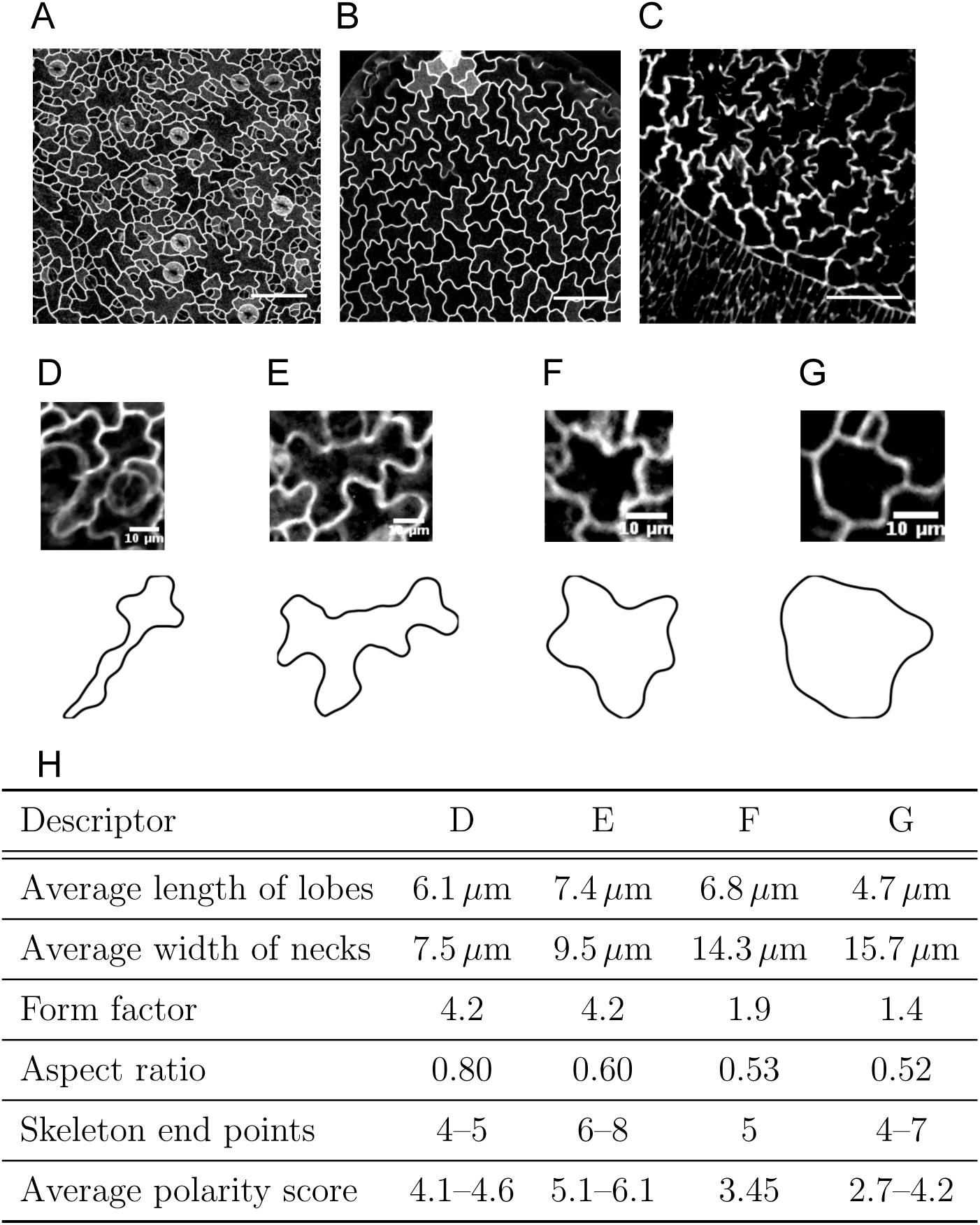
Complex cell shapes and the shortcomings of traditional shape quantifiers. (A–C) Complex cell shapes in both plant (A, B) and animal (C) tissues. (A, B) Pavement cells of *Arabidopsis thaliana* wild type (A) and *speechless* mutant (B), characterised by jigsaw-like shapes consisting of multiple alternating protrusions (lobes) and indentations. (C) Amnioserosa cells in the Drosophila embryo also present cell shapes with similar complexity. Scale bars 50 *μ*m. (D–G) Individual cells from the imaged tissues (upper panel), and the corresponding segmented cell outlines (lower panel). Scale bars 10 *μ*m. (H) Traditional metrics to quantify cell shape can lead to similar values for very different shapes and are sensitive to parameter choices and imaging conditions. Here we compare the cells D–G. See also Figure S1.

Traditional metrics for cell morphology include cell area, cell perimeter, aspect ratio and form factor. Useful as general descriptors, the shape information that can be extracted from them is limited. Very different shapes may yield a similar aspect ratio or form factor (Figure 1D–H). Besides not being unique, such descriptors tend to omit information regarding biologically relevant shape features (Fu et al., 2005). Several approaches classically used to quantify cells, including pavement cells, are summarised in Table 1. Some of those methods, such as the skeleton method, are highly sensitive to image noise as well as to the precise choice of parameters (for an example, see Le et al., 2006). Other metrics, such as lobe length and neck width assessments (Fu et al., 2005), are entirely dependent on humans to judge what a lobe is. Such decisions then strongly impact the average lobe length and neck width measurements (Figure 1 and Figure S1). It renders these metrics highly variable from cell to cell, from phenotype to phenotype and from human to human. In this respect, recently an automatic method was developed to count lobes and indentations (Wu et al., 2016), making this process more reliable. As we will discuss further on, such a method, although already very useful, finds its limitations when the characteristics of the shape reside in the distribution and amplitude of lobes, not only in their number. For instance, some Arabidopsis mutants can present pavement cells with lobes that look more elongated or shallower, but which occur at a similar spatial frequency (Lin et al., 2013).

**Table 1:** Distinct shape descriptors have been used to quantify pavement cells.

| Measure | Description | Reference |
| --- | --- | --- |
| Average lobe length and neck width | Length of each lobe and the distance between opposite indentations within a cell (called necks) are shown in Figure S1. The final measure for a cell is the average of all lobe lengths and the average of all the neck widths. These measurements depend on human criteria to identify lobes and necks, and are given in absolute length units (thus, incomparable throughout growth stages). | (Fu et al., 2005) |
| Form factor (or circularity) | Defined as $\frac{P^2}{4\pi A}$, where $p$ is the perimeter and $A$ is cell area. A circle corresponds to a form factor 1, the lowest value possible. | (Bai et al., 2010; Russ, 2011; Andriankaja et al., 2012) |
| Skeleton | This metric relies on the number of end points of a skeleton representation of the cell shape. The skeleton is formed by iteratively removing pixels from a grid-based cell shape representation, such that eventually a branched one-dimensional graph remains. There are different variants of this algorithm to skeletonise shapes; the resulting branch patterns and length of branches highly depends on the parameters used and are very sensitive to the image resolution. | (Le et al., 2006; Russ, 2011) |
| Average polarity score | Defined as $\frac{c+s}{2}$, where $c$ is the circularity and $s$ the number of skeleton end points. | (Sorek et al., 2011) |

Promising alternatives are methods that take into account the full cell outline information and then simplify the contour of a cell into a series of coefficients that can be employed as shape descriptors in a multivariate study (Ivakov and Persson, 2013; Pincus and Theriot, 2007). One example of these methods is Elliptical Fourier Analysis (EFA), that has been used to quantify two-dimensional complex shapes (Kuhl and Giardina, 1982; Diaz et al., 1989; Schmittbuhl et al., 2003). It takes the coordinates of the outline and decomposes them into a series of related ellipses (described by EFA coefficients) that can be combined to reconstitute that given shape. Despite its wide usage in morphometric studies, EFA cannot retrieve information that directly relates to morphological features of a cell, obstructing biological interpretation. This is because the EFA coefficients are highly dependent on how the cell outline is approximated (a detailed explanation is given in the Supplementary Materials and Methods).

Here we present a new method based on EFA, termed Lobe-Contribution Elliptic Fourier Analysis (LOCO-EFA), that overcomes the common obstacles described above. Our method also uses the information of the whole cell contour but, unlike EFA, provides a set of metrics that directly relate to morphological features, permitting the assessment of cell shape complexity in an objective and automatic manner. Importantly, it is not sensitive to cell orientation or imaging resolution.

To validate the usage of our method on larger cellular datasets, we analyse pavement cell populations from confocal images of *Arabidopsis thaliana*. We then complement this study with the analysis of synthetic tissues generated using the Cellular Potts model (Graner and Glazier, 1992; Glazier and Graner, 1993), in which pavement-like cells have a parametrised specified shape. The latter approach also allows us to touch on a fundamental question in developmental biology, by asking to what degree the resultant cell shape within a confluent tissue context is shaped by cell-to-cell interactions and to what degree it can be explained by intracellular shape control mechanisms. Specifically, here we explore the influence of cell neighbours on the morphogenesis of individual cells by measuring the divergence of their specified cell shape within an isolated context to the shape taken up when being immersed within a tissue. This analysis, performed on *in silico* tissues, allows us to quantify such differences for a generic situation by combining CPM and LOCO-EFA.

## Results

### Quantitative characterisation of cell shape using LOCO-EFA

Applying EFA to quantify cell shapes, we came across a number of specific shortcomings. We first explain those issues to highlight our motivation and choices that led to the development of LOCO-EFA. Detailed mathematical implementation is provided in the Supplementary Materials and Methods. Here, we focus on explaining the analysis in terms of its biological relevance, emphasising how it can be applied and interpreted.

The shape analysis proposed here is linked to harmonic compositions of digitalised cell shape outlines. We find it useful to compare the decomposition of a complex cell shape, such as that of a pavement cell, to the way one decomposes the sounds of musical instruments. This analogy will be used throughout the manuscript for purposes of illustrating the different traits that our morphometric analysis is capturing. For example, when listening to a note generated by an instrument, a quantifiable observable is the pitch. Within the context of pavement cell shape this corresponds to the observed number of lobes or, as we will explain in detail, to the dominant spatial frequency of the pavement cell outline. Another quantifiable property of musical note is its volume, the amplitude. In terms of cell shape this corresponds to the extent to which lobes and indentations are protruded and retracted. Finally, the timbre of musical instruments is what essentially distinguishes, for example, a clarinet from an oboe, as both play the same note (same pitch) at the same volume. An analogous notion in cell shape studies would be the ability to capture additional aspects of cell shape morphology, as well as to recognise subtle differences between cells, even when the number of lobes and their level of protrusion is the same.

As a starting point, Elliptical Fourier Analysis (EFA, Kuhl and Giardina, 1982) can be used to describe the contour of any complex two-dimensional shape, including non-holomorphic shapes such as pavement cells, which most other methods are unable to capture (see Figure S2 and Supplementary Materials and Methods). Using the coordinates of the two-dimensional (cell) outline (Figure S3A), EFA decomposes the shape into an infinite series of ellipses (also referred to as “modes” or “harmonics”). This series of ellipses, *n* = 1 *… ∞*, can then be combined in the following way to exactly retrieve the original shape: each *n*th elliptic harmonic traces *n* revolutions around the first ellipse while orbiting around the previous (*n* − 1) harmonic ellipse, which in its turn is orbiting around its previous one (*n* − 2), and so forth (Figure S3B). This summation results in an outline being ‘drawn’, as is visualised in Movie S1. Obviously, a cut-off has to be chosen for the number of modes that are taken into account. In general this will be at a value for which the cell contour description is reasonably close to the original cell outline. The fact that each ellipse represents a certain harmonic suggests that it captures dominant spatial frequencies within the original shape. This is why it has been considered to be a reasonable descriptor for cell outline properties (Schmittbuhl et al., 2003).

When shapes are quantified using the EFA method, however, the pitch, i.e., the most basic cellular feature to quantify, is actually not directly retrieved, not even for the most simple shapes. For instance, one might expect a six-sided shape to present a strong contribution from the 6th mode. Instead, the EFA method represents such a shape as a mixed contribution from the two adjacent modes, the 5th and 7th mode (Figure 2D and Figure S3C). This mismatch is a consequence of how the individual ellipses obtained from decomposing the outline contribute to the outline. Each ellipse does so by rotating either clockwise or counterclockwise. The direction of this rotation in respect to the direction of rotation of the first mode causes either an increase or a decrease in the actual number of features being drawn, always one off from the actual mode (Figure S3D, E, Movie S2 and Movie S3). As a consequence, the ‘pitch’ obtained using EFA does not correspond to actual cell features, which hinders interpreting the resultant values of this method. Moreover, EFA coefficients on their own are redundant, i.e., there are more parameters than needed to specify the same specific shape (Haines and Crampton, 2000). Therefore, comparison of cell shapes on the basis of their EFA coefficients (for example, by means of principle component analysis) becomes nonsensical. Together, these traits make the EFA method unsuitable for reliable quantification of cell morphology and renders prohibitive any meaningful comparisons between multiple cell outlines.

**Figure 2:**
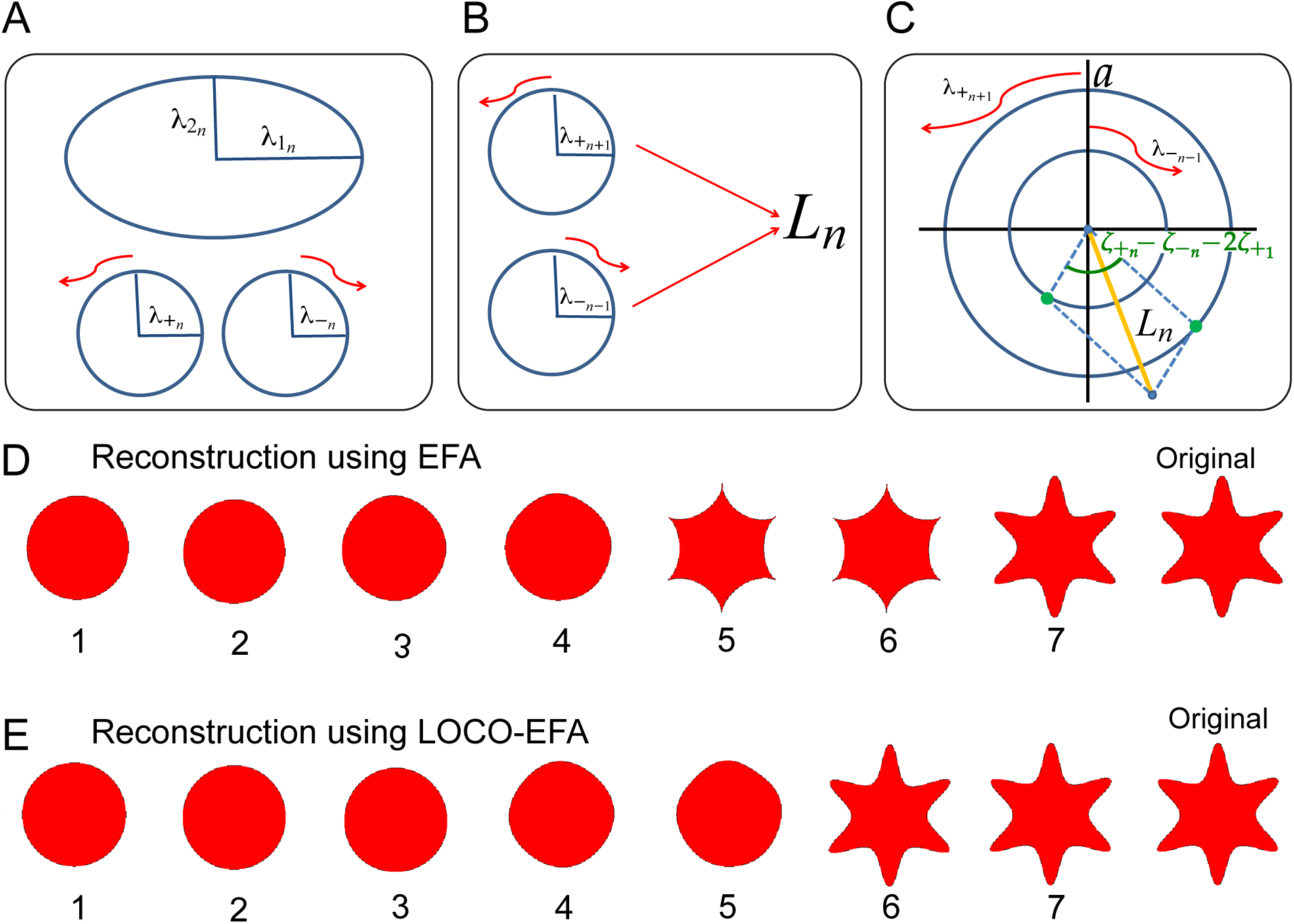
To get LOCO. (A) Each EFA elliptic harmonic is decomposed into two counter-rotating circles. (B) Mode *L*_*n*_ is composed of the counter-clockwise rotating *n* + 1th EFA harmonic circle and the clockwise rotating *n* − 1th circle. (C) The combined amplitude contribution to *L*_*n*_ (yellow line) of the two counter-rotating circles with radii 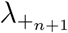 and 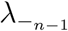 is also dependent on the offset in their starting points and the offset of the overall (mode 1) starting point, which together determine the initial phase shift (green dots) in the amplitude contribution of each rotor. (D, E) Comparison of closed contour reconstruction through either EFA (D) or LOCO-EFA (E). Although both approximations eventually converge to the original 6-lobed star shape (labelled ‘Original’), the reconstruction using EFA harmonics (D) generates a spurious shape after addition of the 5th harmonic and only recovers the original shape after addition of the 7th harmonic, whereas the reconstruction using LOCO-EFA (E) reconstitutes the original shape only at the 6th mode, matching the number of protrusions. The number of modes used for each sequential reconstruction is indicated below each shape.

Diaz et al. (1990) proposed a heuristic solution for the mismatch between actual shape features and EFA’s dominant harmonics, using the fact that the relative direction of rotation is a main determinant of the generated pitch (see Supplementary Materials and Methods for details). It turns out, however, that each ellipse is simultaneously contributing to two different spatial frequencies, something this heuristic can not solve (see Figure S3F and Movie S4). As a consequence, although their method is often (but not always) able to correctly recapitulate the ‘pitch’, it is never able to correctly capture the amplitude or timbre of the cell shape. Further details regarding the range of issues when using EFA are discussed in depth in the Supplementary Materials and Methods.

To overcome these limitations we essentially propose a new basis for the outline reconstruction, which we have coined *ℒ*_*n*_, after lobe number. In a similar manner as EFA, the modes can be summed up to recreate the original shape and each individual mode is captured by a set of four unique parameters. However, there are two important distinctions. Firstly, any possible cell outline is now only represented by one unique combination of *ℒ*_*n*_ coefficients. In the Supplementary Materials and Methods we discuss in detail the steps required to eliminate all levels of redundancy for shape analysis. Secondly, cell shape features such as the protrusion number (the “pitch”), their amplitude (the “volume”) and the characteristic lobe distributions (“timbre”) are directly represented by the *ℒ*_*n*_ coefficients. The *ℒ*_*n*_ coefficients are obtained after decomposing each EFA harmonic into its exact, specific contributions to two separate *ℒ*_*n*_ modes (Figure 2A–C). In general, the EFA modes *n* − 1 and *n* + 1 both partly contribute to mode *ℒ*_*n*_, although some specific exceptions apply (see Figure S4). The resulting method, which we coin Lobe Contribution-EFA or LOCO-EFA, thus consists of: eliminating multiple representations of a given outline; decomposing each *n*th elliptical EFA mode into two separate lobe contributions; and finally integrating those separate modes into single LOCO-EFA modes. Every *ℒ*_*n*_ mode can then be regarded to represent two oppositely rotating circles, each with its own starting point for the rotation, the four *ℒ*_*n*_ coefficients providing the radii and starting angles of both circles. We next assign a scalar *L*_*n*_ value, to specifically capture the amplitude contribution (“volume”) of each mode. We found that in order to correctly capture this, both the radii of and the angular distance between the starting points of the two contributing circles, as well as the starting point of the main circle, *ℒ*_1_, have to be taken into account (see Figure 2C, and Supplementary Materials and Methods for further details regarding its derivation). Calculating the *L*_*n*_ values results in a spectrum representing the relative contribution of each individual mode to the cell shape (Figure S3C). For instance, the spectrum of the six-lobed test shape used for Figure S3C indeed contains a pronounced peak at mode six, as well as a peak at mode one, which represents the overall circular shape of that outline. The original shape can be reconstructed using all modes up to a specific mode number. This allows us to visually appreciate the specific contribution of mode six to the six-lobed test shape (compare Figure 2E with Figure 2D).

To further illustrate how LOCO-EFA characterises shapes, we apply it to simple geometrical shapes with variable numbers of protrusions (Figure 3A–I). Indeed, LOCO-EFA robustly determines the main *L*_*n*_-mode of each shape corresponding to the correct number of lobes (Figure 3J). Each *L*_*n*_ profile provides information about the composition of all morphological periodicities contributing to the shape, reflecting the geometry being considered. We next tested if LOCO-EFA also correctly captures the “volume”, by applying the method to generated shapes of the same “pitch” (i.e., having a same number of lobes), but with variable amplitudes (Figure 3N–Q). Indeed, the magnitude of the *L*_*n*_ components changes accordingly (Figure 3T), and their absolute value quantitatively corresponds to the size of the extensions. Thus, LOCO-EFA not only retrieves the main number of morphological features of hypothetical cell shapes, but also a range of additional characteristics.

**Figure 3:**
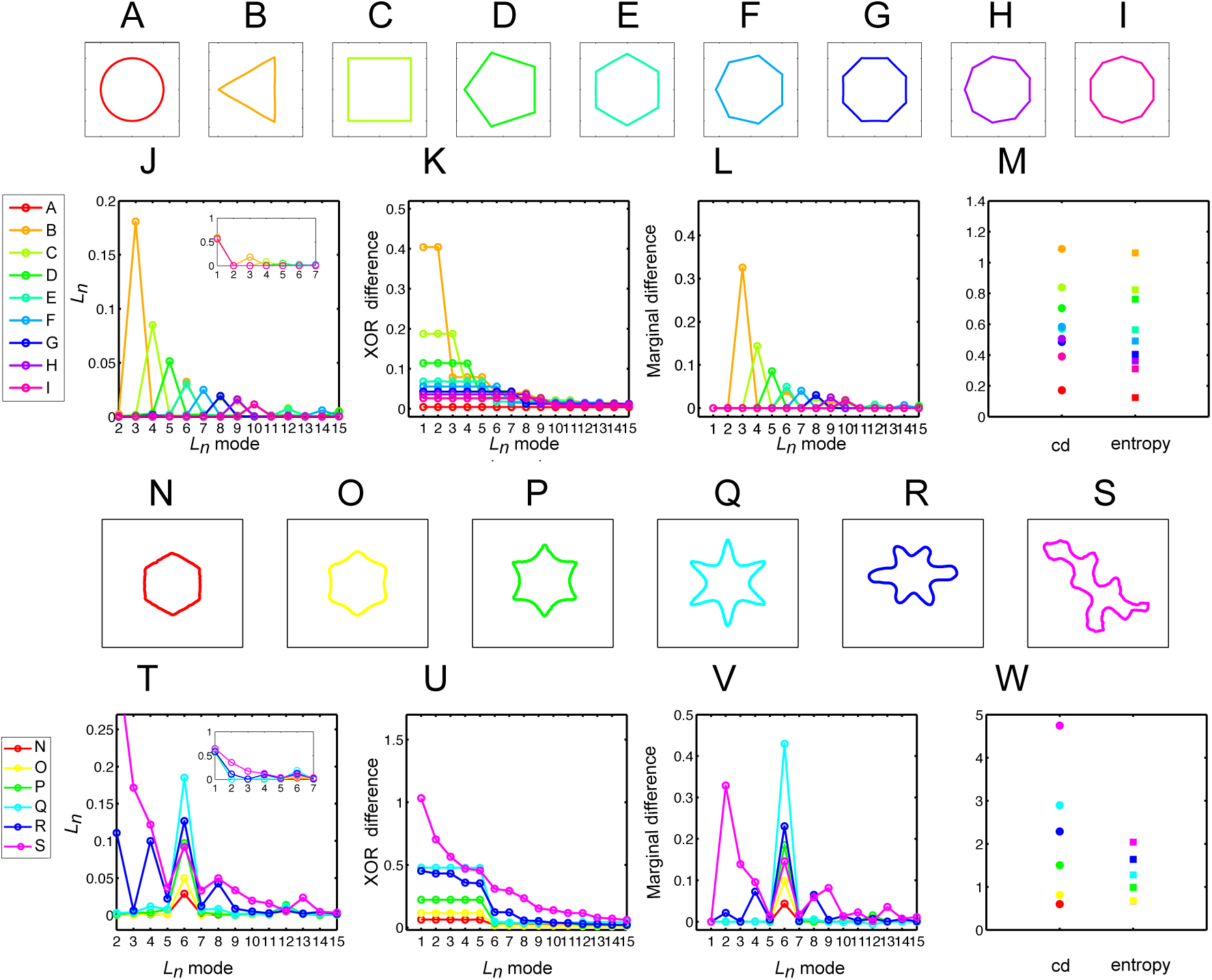
Interpreting LOCO-EFA-derived measures for geometrical and asymmetric shapes. (A–I) Symmetrical and well-defined geometrical shapes with normalised area. (J–L) *L*_*n*_, XOR, and marginal difference profiles for shapes (A–I). (J, L) For each geometric shape, a clear peak appears in the profiles, this main contributor to the shape always coinciding with the number of protrusions. (M) Cumulative difference (cd) and entropy for shapes (A–I). (N–Q) Symmetrical shapes with increasing protrusion amplitude. (R, S) Asymmetrical shapes. (T–V) *L*_*n*_, XOR, and marginal difference profiles for shapes (N–S). Increasing protrusion amplitude leads to increasing peak levels in the profiles. Asymmetric shapes present multiple peaks, indicating that multiple modes are needed to recapitulate the original shape. (W) Cumulative difference (cd) and entropy for shapes (N–S).

Following the analogy of sound decomposition, a more nuanced perception for quantification is timbre. Timbre can be understood as residing in the additional amplitude spectrum. It is determined by which overtones are emphasised in relation to one another. For cell shape studies, we consider “timbre” analysis the ability to capture additional aspects of the cell shape complexity, besides the main number of protrusions/lobes which a cell presents. This additional information should enable distinction between different cellular phenotypes, such as between wild type and mutants (Lin et al., 2013). To illustrate, Figure 3R, S show two additional six-sided shapes which differ in “timbre” from Figure 3Q, with their accompanying *L*_*n*_ values (Figure 3T). For both shapes, there is a clear peak at *L*_6_, reflecting their six-lobedness, and additional peaks or overtones, such as *L*_2_, capturing the elongated nature of these shapes, and so forth. Thus, LOCO-EFA retrieves not only the main number of morphological features of an hypothetical cell (i.e., lobes), but also important fine-grained characteristics.

From the full characterisation of cell shape by *ℒ*_*n*_ modes, additional objective metrics can be derived to help quantify different aspects of“cell shape complexity”. We here define four metrics: XOR difference; marginal difference; cumulative difference; and entropy.

Firstly, complementary to the magnitude of each individual *L*_*n*_ mode, the cell shape complexity can be estimated using the information of the approximation to the original shape by the first *N* LOCO-EFA modes only. It addresses the question how relevant each subsequent *L*_*n*_ mode is for explaining that specific shape. Figure 2E already illustrated the importance of a specific mode for reconstructing the original shape (in that case, mode six). One can straightforwardly quantify the relative contribution of that mode for explaining the shape by calculating the total difference (in area, expressed as either number of grid points or *μ*m^2^) between the original shape and the superimposed reconstructed shape when the first *N* LOCO-EFA modes are used. To do so, we take the *XOR* (exclusive or) between the original and reconstructed cells (see Figure S5). In this context, a more “complex” shape is one in which more LOCO-EFA modes are needed to obtain a good match between the reconstruction and the real shape (i.e., when the *XOR* approaches zero only at higher *N* numbers). For this criterion, note that a circular cell can be perfectly reconstituted using only the contribution of the first LOCO-EFA mode (*N* = 1), and is therefore the least complex shape with an XOR value of zero for the whole spectrum. On the other hand, cells presenting a high number of heterogeneous lobes require a high number of modes for *XOR* to approach zero (Figure 3). This can be observed in Figure 3K, U, which show the *XOR* difference profiles for the series of test shapes.

This manner of defining cellular complexity can be further compressed by taking the total area under the *XOR* difference curve, a scalar quantity which we call the cumulative difference (cd). It derives a single value from the XOR spectrum, with higher values for more complex shaped cells. The more similar the cell shape is to a circle (which can be perfectly described using the first LOCO-EFA mode only), the more the cd value approximates zero. Figure 3M, W shows the cd values for the series of test shapes. From these, it becomes clear that cd becomes high for shapes that strongly deviate from circular, it is therefore not just correlated with having many lobes. In general, however, when morphological protrusions increase in number or become larger in amplitude, the cumulative difference also increases since higher order modes are required to approximate the shape.

*XOR* profiles typically do not change smoothly. Instead, some modes peak as they strongly contribute in capturing the main features of the shape. Hence, the marginal decrease in the *XOR* value when an extra mode is added, coined *marginal difference*, provides additional information about the shape’s dominant modes (Figure 3L, V). This measure is comparable to the *L*_*n*_ values, also determining something akin to “pitch” and “amplitude”. We found, however, that it bears a higher discriminatory power for more complex and irregular cell shapes. Moreover, in the case that a cell shape has significant contributions from a combination of different modes, then high marginal difference levels can be directly linked to specific cellular features (see Figure S5). Thus, the marginal difference can help identifying which modes are the most relevant for specific shape aspects.

Finally, one can argue that shape complexity is not solely about high numbers and large amplitudes of the protrusions, but instead reflects overall irregularity of the protrusions. For example, with the previous measures a highly regular star-shaped cell with five outspoken lobes could be quantified as being as complex as a highly distorted cell with different amplitudes and distributions of five lobes, albeit less pronounced than the starshaped case. One might therefore prefer to define cell shape complexity as the tendency of a cell to deviate from geometrically well-defined periodic outlines. A useful measure for this alternative definition of “cell shape complexity” is calculating the Shannon entropy *E* of the *L*_*n*_ spectrum. The *entropy* measure is based upon the information content contained within the distribution over the whole *L*_*n*_ profile (see Equation 47 in the Supplementary Materials and Methods). For many shapes, the entropy measure yields very similar results to the cumulative difference. They diverge for cell outlines having a strong contribution from the lower modes. We observe that in these cases the entropy values deliver more meaningful results in regard to what one considers being more “complex”. This is due to the fact that lower modes can have a large contribution to the cumulative difference. For example, there can be a high contribution from *L*_2_ in the case that a cell is very elongated. Simply being elongated, however, does not so much represent shape complexity in the way defined above. For such a simple elongated shape, the cumulative difference (i.e., the integral or area under the curve of the *XOR* between the original and reconstructed profile) can be very similar to a cell with contributions distributed among many modes. We would consider the latter outline, however, to be much more ‘complex’. The entropy measure is able to correctly capture this form of complexity.

Next, we measured the *L*_*n*_ profiles and corresponding cell shape complexity measures for a population of real cell shapes, which are much less geometric than the idealised cases treated so far.

## LOCO-EFA applied to plant pavement cells

To validate our method, and its derived measures introduced above, we analysed *Arabidopsis thaliana* leaf epidermal pavement cells, individual cells that were tracked *in vivo* over time as well as whole populations of cells at a single time point. Actual biological cells, such as pavement cells, can be highly asymmetrical, with their *L*_*n*_ landscape characterised by multiple peaks (Figure 3S, T). This is because the outline of an asymmetrical cell with a given number of protrusions placed quasi-periodically along its edge can be interpreted as multiple protrusion frequencies superimposed. In general, the total number of hand-counted lobes in a cell image matches to a peak at the corresponding *L*_*n*_ value (but note here that hand-counting is very subjective). For instance, for nine lobes a peak will be observed at *L*_9_. However, if the lobes are distributed in a more or less pentagonal clustered fashion, this would lead to an additional peak at *L*_5_, whilst superimposed on a triangular shaped cell basis a *L*_3_ contribution would be found, and so forth.

Pavement cells acquire their characteristic jigsaw puzzle-like shape through multipolar growth patterns, such that relative simple shaped cells become highly complex over time (Figure 4A–G). Notably, the smooth shape changes over time are clearly reflected in the *L*_*n*_ profiles over time (Figure 4I). Its initial squarish shape and later nineand thirteenlobeness are well captured by the LOCO-EFA method, through peaks at mode *L*_4_, *L*_9_, and *L*_13_, and corresponding peaks in the marginal difference profile. In contrast, when the EFA method is used (Kuhl and Giardina, 1982), the third and fifth mode are erroneously indicated to represent shape features, as well as other mismatches (Figure 4H). Importantly, the smooth transition in cell shape development over time is captured by a corresponding smooth change in the LOCO-EFA *L*_*n*_ profile for the different time points, but by highly irregular changes in the EFA profile. This illustrates that very comparable shaped cells have very different EFA profiles, making EFA further unsuitable to analyse real pavement cell populations. Figure S6 presents the robust time dynamics analysis for yet another pavement cell using LOCO-EFA.

**Figure 4:**
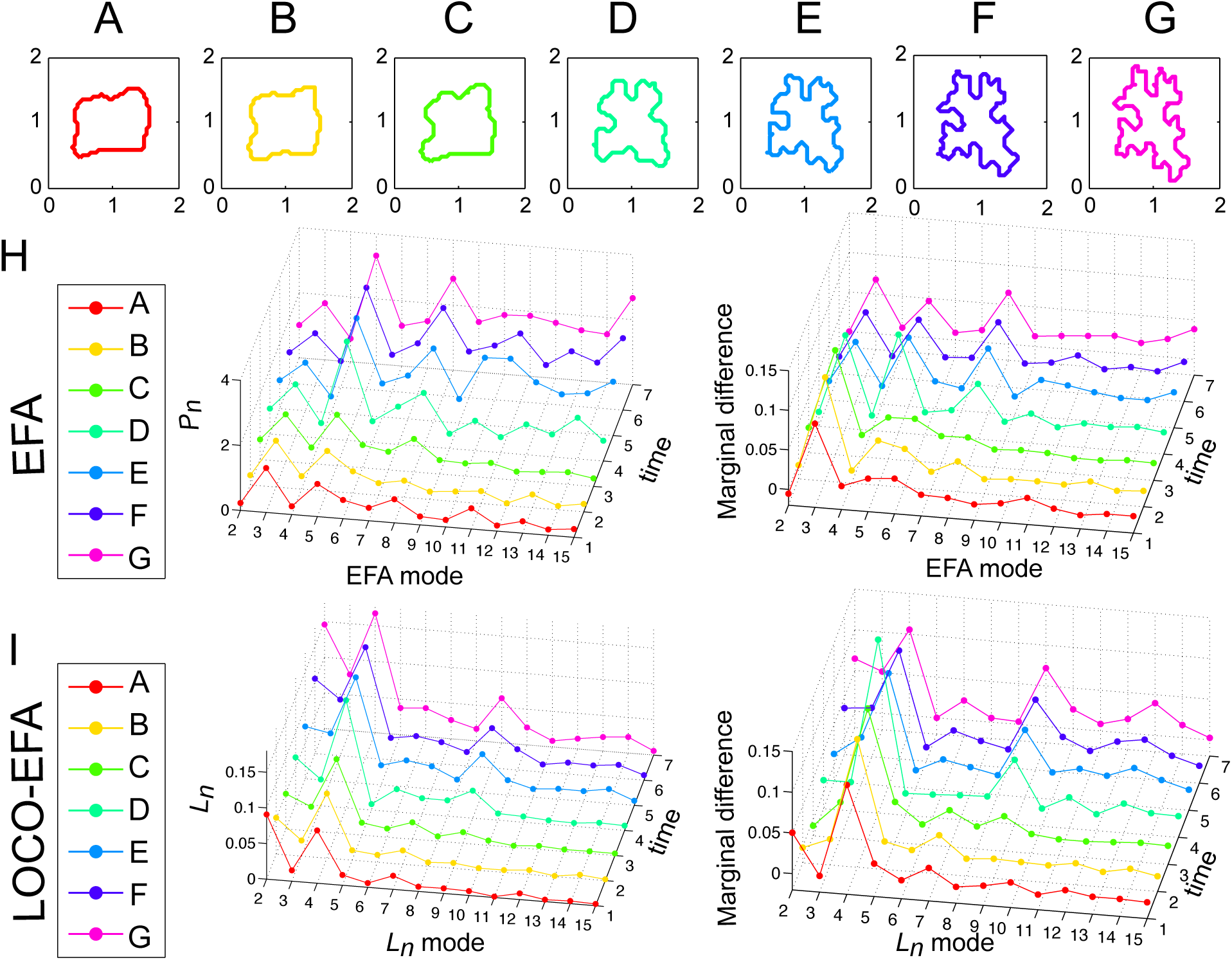
LOCO-EFA metrics on a cell changing its shape over time. (A–G) Sequence of a tracked pavement cell growing over time with normalised area. (H) *P*_*n*_ and marginal difference profiles using EFA. Applying EFA modes to approximate the cell shapes leads to erratic profiles that fail to recover the biological sequence of development, as can be observed in the *P*_*n*_ profile and as spurious peaks at the third and fifth harmonics in the marginal difference profile. (I) *L*_*n*_ and marginal difference profiles using LOCO-EFA. The LOCO-EFA measurements recover the smooth transitions during the cell morphogenesis. The overall square symmetry of the cell is captured by a peak at *L*_4_, the formation of lobes by a smooth increase in *L*_9_, and later *L*_13_.

To visualise the shape characteristics of populations of pavement cells, we applied the analysis to leaves of the *speechless* mutant (MacAlister et al., 2007), which does not generate during the leaf development any other cell types such as meristemoids or stomata (Figure 1B, Figure 5A), as well as to wild type leaf epidermis, consisting of pavement cells, stomata and other cells from the stomatal lineage (Figure 1A, Figure 5B).

**Figure 5:**
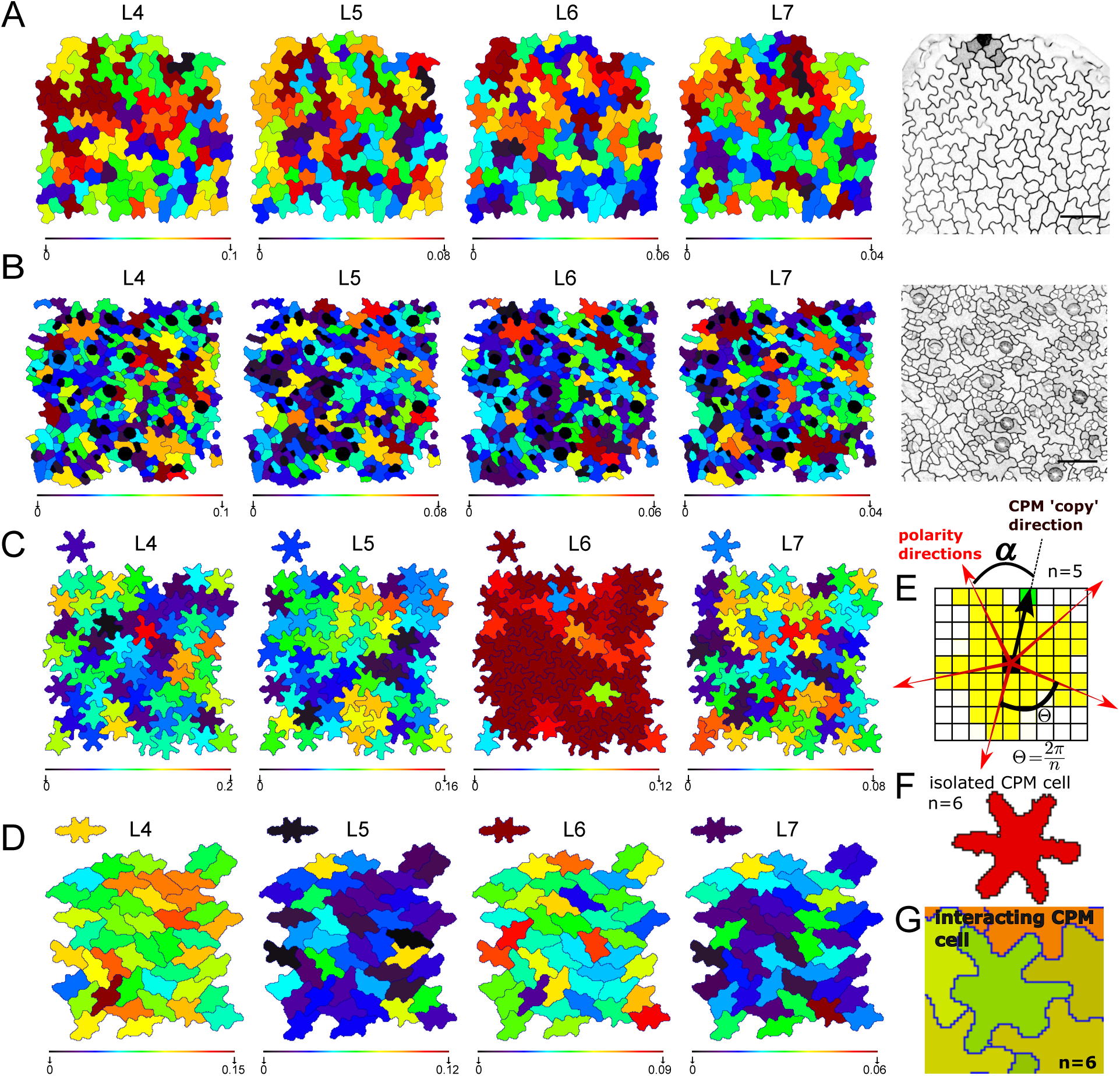
LOCO-EFA analysis on *in vivo* and *in silico* pavement cells. (A, B) LOCO-EFA applied to *speechless* mutant (A) and wild type (B) leaf tissue. Colour coding depicts the *L*_*n*_ values for 4 different *L*_*n*_ modes (*n* = 4 *…* 7), as indicated above each panel, with the scale shown below each panel. It shows that very few cell shapes can be reasonably captured through a single *L*_*n*_ mode, revealing the complexity of the cell shapes. (C, D) LOCO-EFA applied to *in silico* pavement cells reveals the degree of divergence from their specified shape that interacting cells within a tissue experience. Two different specified cell shape populations are shown (SCS1 and SCS3, each with six lobes, see Table S1). The specified shapes are depicted above each panel. Colour coding within the panels and of the specified shapes above each panel again depicts the *L*_*n*_ values, with the scale shown below each panel. Because cells interact with each other within the tissue, strong deviations in *L*_*n*_ contributions can be observed. (E–G) Modelling framework to generate the *in silico* tissues. (E) Standard Cellular Potts Model is modified to allow for a specified number of lobes (here, *n* = 5) to be formed at a regular radial spacing (*α*). (F) This gives rise to a symmetric, multi-lobed specified cell shape, shown in red. (G) Within a tissue context, however, the same specified shape deforms as to accompany the neighbouring cells.

Using LOCO-EFA, it is straightforwardly possible to dissect the precise contribution of each mode for each cell in the population. Figure 5A, B show the spatial distribution of cells within a tissue which are predominantly 4-, 5-, 6-, or 7-lobed, by colour-coding the cells according to those specific *L*_*n*_ values. Very few cell shapes present only a single high *L*_*n*_ value. Instead the majority of shapes are composed of significant contributions stemming from multiple modes. Consequently, simply manually counting, or applying automatic counting algorithms, for such biological cells the number of lobes would lead to incomplete information regarding their shape. It would be difficult to compare mutant phenotypes. Moreover, our data shows pavement cells do not have a population-wide preferential *L*_*n*_ with a highest contribution to the shape (Figure 5A, B).

The heterogeneity in modes that composes real populations of pavement cells suggests that their resultant cell shapes cannot be explained solely by currently proposed intracellular molecular mechanisms underlying lobe and indentation patterning. This is because those mechanisms, based on the existence of two counteracting pathways (one for lobe formation and another for indentation formation, see details in Xu et al., 2010) should give rise to Turing-like instabilities which are therefore expected to generate symmetrical shapes (Vanag and Epstein, 2009). Moreover, these models would predict that equally sized cells exhibit equal lobe numbers. However, the cell shape patterning takes place within a confluent tissue, which obviously complicates how individual cells generate their shape. In the experimental setting it is very hard to distinguish between the preferred shape of each individual cell due to its intracellular patterning, and the constraints imposed by the tissue. It is well-known that if cells prefer to be round, they will take up an hexagonal shape within a tissue context (Thompson, 1917), but it is unclear what to expect for multi-lobed shapes. Therefore, to both better understand what should be our expectation when a population of cells with complex shape preferences form a confluent tissue alike the leaf, and to further validate the LOCO-EFA method on populations of cells, we performed cell-based simulations of interacting cells with prespecified cell shape preferences, and employed LOCO-EFA on the resulting *in silico* tissue.

### **Applying LOCO-EFA to *in silico* populations and the effect of interactions between preferred cell shapes**

We create *in silico* cells using the Cellular Potts Model (CPM), an energy-based framework that allows us to represent cells and their dynamics through small membrane extensions and retractions (see Methods section). In its basic form, CPM cell shapes emerge due to the interaction between interfacial tension (the strength of which can depend on the cell types involved), internal cellular pressure and cortical tension (Magno et al., 2015). Here we used an extension of the CPM which predefines intrinsic forces causing elongation and lobeness, resulting in more complex cell shapes. This extension consists in applying additional, cell-specific forces each time a cell extension or retraction is considered, giving rise to elongated and/or multilobed preferred cell shapes (Equation 3). Three additional forces are used that capture (i) an intrinsic tendency to elongate; (ii) a tendency to form a specified number of lobes; and (iii) an additional force for the cell to round up (Figure 5E–G, van Rooij et al. (2017a) and Movie S5). The latter term robustly prevents cells from falling apart, especially within a confluent tissue with many conflicting preferred cell shapes. A population of cells, all with a same preferred shape, were then allowed to interact with each other within a tissue context. In this way we can compare the shape of a single cell in isolation with the shape cells attain within a tissue.

We here present the analysis on two distinct specified shapes (Figure 5C, D; see Table S1 for the specific parameters used). Both preferred shapes have six lobes, the cells shown in Figure 5D moreover tend to be elongated. Above the panels we plot the shape that the cells attain in isolation. Although the same cell shape is specified for all cells within the population, cells acquire very different shapes due to their local interactions. We quantified this diversity using LOCO-EFA. This was done by colour-coding the contribution of *L*_4_–*L*_7_ (as indicated for each panel), for both the isolated cells and for the resultant shapes of all the cells within the simulated tissue. As expected, the isolated cells present a very high contribution by *L*_6_, with marginal contributions from the other modes. In contrast, due to the cellular interactions within the tissue, other modes can also become prominent, highly varying from cell to cell, even though each cell has exactly the same specified shape. Thus, although a cell in isolation would be able to generate regular protrusions with specific amplitudes, the periodical lobe formation becomes inhibited in a packed tissue environment, with symmetry and shape distortions being a direct consequence of tissue packing (Figure 5C, D). Such a tendency was observed for all simulations performed, irrespective of the specified cell shape, i.e., the number of lobes, their amplitude, and the level of overall cell elongation, as well as over a wide range of parameter values for the cell-cell interactions (Figure S7, Figure S8). Radially, well-spaced periodically lobed cell shapes are not likely space filling, hence resolving the competition between preferred shape and confluency seems to be a universal driving force to complex cell shapes.

## Discussion

The progress in microscopy and imaging techniques within the biological sciences generates a need for adequate analytic tools to capture relevant information efficiently and objectively (Zhong et al., 2012). Image acquisition through high-throughput microscopy approaches generates amounts of data that are beyond the human ability (or patience) to be analysed manually, demanding computational automatic tools. We have developed a new analytic tool which takes as the input the contour of a 2D cell projection, and extracts from it, in an efficient and parameter-independent manner, quantitative meaningful cell shape information. Importantly, the pipeline can be integrated within other recent advances in segmentation techniques (Fernandez et al., 2010; van Rooij et al., 2017b), to fully automate shape analysis of series of images.

Our method can intuitively be understood using the analogy of music perception. To quantify an instrument playing a certain note, say a violin playing the note A, one would firstly wish to have a device that can capture which note is played. We have indeed shown that LOCO-EFA, unlike EFA, is able to correctly determine the analogous feature for shapes, which is the number of protrusions.

A recent method developed by Wu et al. (2016) can also be employed to count the number of protrusions. When the biological question asked requires not only the “pitch” to be measured, but also the “volume” and “timbre”, corresponding to lobe amplitude and other irregularities, such a method will be insufficient. Indeed, LOCO-EFA provides a broader spectrum of the shape properties, a holistic set of measurements that allows complex morphologies to be quantified in a reproducible manner.

We illustrated how the measurements obtained with LOCO-EFA can be interpreted, first using simple shapes (geometrical or symmetrical forms), then extending the analysis to highly asymmetric shapes, for which the pavement cells of Arabidopsis are a good paradigm. To assess the performance of our technique, we extended our study to confocal microscopy images of populations of pavement cells. Such a shape analysis is biologically relevant, because many of the players accounting for the lobe and indentation patterning are known (Xu et al., 2010; Grieneisen et al., 2017), enabling one to extend the study of cell shape control to mutants and experimental interferences in the future. We found that very few cells have a symmetrical shape, i.e., they can not be represented well by a single high *L*_*n*_ value. Such composition in real cell shapes of several *L*_*n*_ values is not likely to be explained only by the existence of two counteracting pathways specifying lobe and indentation identities. Our *in silico* approach instead suggests that the dynamics of space-filling complex shapes can dramatically increase the overall irregularity: even when the CPM cells are specifically programmed to develop well-defined characteristic shapes, the interactions between them trigger the formation of highly deviating cell shape characteristics. As a result of being within the tissue, the main, specified mode decreases in strength while other modes become relevant. Altogether it leads to very asymmetrical resultant shapes, even though very symmetrical shapes were specified.

Although our synthetic data is but a phenomenological reconstruction of real shapes, our results suggest that the local influence of neighbours during pavement cell development should be important for the acquisition of their final shape. To assess this hypothesis in further studies it will be crucial to perform quantitative shape analysis on *in vivo* cell populations over time, combined with growth tensor analysis (i.e., determination of the spatial pattern in growth rate and its anisotropy). Such studies, combined with genetic or physical perturbations in cell growth and deformation, and with *in silico* cell growth models, should allow untangling of the relationship between the cell shape as specified at the cellular level and the resultant shape arising from the complex interactions between the cells at the tissue level.

When LOCO-EFA was applied to cell tracking data, we observed that the profiles of those dynamically changing cells varied smoothly over time. Such trajectories are distinct from cell-to-cell and provide unique fingerprints of each individual developing cell. This opens the possibility of using the *L*_*n*_ modes as cell identifiers within a temporal sequence of images, to help track populations of cells automatically.

Although we have here applied LOCO-EFA to objectively measure cell shape differences in pavement cells, providing us a powerful tool to evaluate pavement cell shape dynamics and compare plant phenotypes, it can also be applied to different cell types with complex cell shapes of any species, as well as to any other layouts, such as entire leaves, within or outside the realm of biology. Recently, Fourier analysis was applied to quantify myosin polarity (asymmetrical distribution of molecules) in animal epithelia (Tetley et al., 2016). In a similar way, the LOCO-EFA method could be applied to quantify the cellular distribution of molecules within complex cell shapes. Finally, our method could be integrated within recent image analysis pipelines, allowing one to extract and analyse cell shape information in a high throughput manner (van Rooij et al., 2017b; Heller et al., 2016; Stegmaier et al., 2016).

## Material and Methods

### Confocal images and image processing

Columbia wild type or *speechless* mutant (MacAlister et al., 2007) leaves expressing pmCherry-Aquaporin (Nelson et al., 2007) were imaged using a confocal microscope Leica SP5 at comparable stages and in comparable regions. Further image processing was done using ImageJ and images were segmented using in-house software (Segmentation Potts Model (van Rooij et al., 2017b)). Cells changing over time were imaged using a custom-made perfusion chamber (Robinson et al., 2011; Sauret-Güeto et al., 2012; Kuchen et al., 2012).

### Shape descriptors

Average lobe lengths and neck widths were calculated using ImageJ (Analyse *→* Measure). The skeleton was calculated using Better Skeletonization by Nicholas Howe, available through MATLAB File Exchange.

### Geometric shapes

All geometric shapes were generated by the “superformula” described in Gielis (2003), and were analysed in the same manner as the confocal images.

### XOR and other measurements

All the grid points belonging to each individual real or synthetic pavement cell were compared to all the grid points captured by the subsequent series of LOCO-EFA reconstructions. A reconstruction of level *N* takes into account the first *N ℒ*_*n*_ modes. The *in silico* cells were generated using the Cellular Potts Model, which is a grid-based formalism, while for the experimental data the grid points were directly defined by the imaging resolution. The scripts used to calculate the XOR and to colour-code the real and synthetic cells were written in the coding language C. Shape approximations, cumulative difference and entropy were calculated using the first 50 *L*_*n*_ modes. To capture cell shape complexity linked to protrusions rather than mere anisotropy, cumulative difference is calculated from the second *L*_*n*_ mode onwards. This value turned out to be more than sufficient to capture any cell shape given the grid point resolution used for all cases here. Note, however, that very high resolution images might require additional modes to fully capture the shape.

### Cellular Potts Model generating complex cell shapes

The Cellular Potts Model (CPM) is an energy-based model formalism used to model cellular dynamics in terms of Cell Surface Mechanics (Magno et al., 2015). Individual cells are described by a set of grid points on a lattice. In this manuscript we used the CPM to generate *in silico* cells with relatively complex shape preferences that are allowed to interact within a confluent setting. During each simulation step a grid point is chosen in a random fashion to evaluate whether its state changes into one of its neighbouring states, effectively corresponding to a small cell shape modification at that point. To evaluate if such state change will occur, the energy change is calculated that such a copy would cause. This is done by calculating the change in the configurational energy as defined by the following Hamiltonian, which sums up the energy contribution of each pixel within the entire field as well as of all cells:

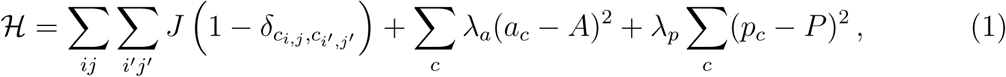

*J* refers to the coupling energy, summed over all grid points (*i, j*) and their eight (2nd order) neighbours (*i′, j′*). The Kronecker delta term 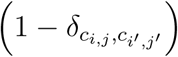 simply assures that neighbouring lattice sites of the same state (i.e., belonging to the same cell) do not contribute to the total energy of the system. The variables *a*_*c*_ and *p*_*c*_ denote respectively the actual cell area and the actual cell perimeter for each cell (*c*); the parameters *A* and *P* denote the target cell area and perimeter. The parameters *λ*_*a*_ and *λ*_*p*_ describe the resistance to deviate from the target area and perimeter, respectively. The probability a copying event is accepted depends on the change in the Hamiltonian, *Δℋ* = *ℋ*_*after*_ - *ℋ*_*before*_, in the following way:

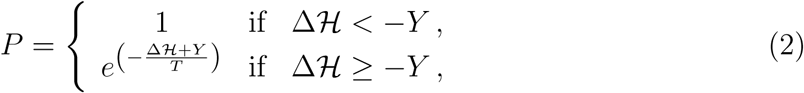

where *Y* corresponds to the yield or ability of a membrane to resist a force and *T* (simulation temperature) captures additional stochastic fluctuations. Copying events which decrease *ℋ* by at least *Y* will always be accepted, otherwise acceptance follows a Boltzmann probability distribution (Equation 2).

To generate cells with a particular number of preferred protrusions, we modify the change in the Hamiltonian as is calculated for every evaluated copying event, effectively shortcutting intracellular biochemistry and biophysics, in the following way. Simulated cells are attributed with a specified preferred number of lobes, amplitude of lobes, overall elongation and roundness, implemented by modifying the change in the Hamiltonian for every evaluated copy event as follows (van Rooij et al., 2017a):

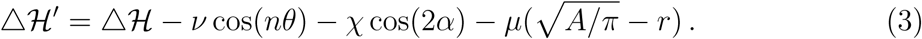

Those three additional terms are evaluated for both cells involved in the copying event, so there are effectively six additional terms. The first term captures the tendency to form *n* lobes, with *v* capturing the propensity to extend to form a lobe or to retract to form an indentation, thus giving rise to the amplitude or pointedness of the lobes. *θ* describes the angle between any of the *n* equally spread out target directions for outgrowth and the vector determined by the coordinates of the grid point under evaluation and the centre-of-mass of the cell (hereafter called the copy vector) (Figure 5E). To clarify, when a cell extension is considered right on top of one of the target directions, then *nθ* = 0, cos(*nθ*) = 1, and tendency to extend is maximally increased, while halfway two target directions, *nθ* = *π*, cos(*nθ*) = −1, and the tendency to extend is maximally suppressed.

The second term in Equation 3 captures an overall elongation, implemented in a similar fashion. The parameter *χ* corresponds to the propensity to elongate and *α* is the angle between the elongation vector and the copy vector.

If only these two terms are used, cells within tissue simulations can easily lose coherence, i.e., fall apart. Therefore a third term was added, capturing a propensity to roundness. The parameter *μ* captures the resistance of a cell to deviate from a circle, with *r* being the length of the copy vector, and 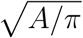 being the preferred radius of cell, given its target area. For further details, see van Rooij et al. (2017a).

Importantly, the target lobe and elongation vectors are not fixed during the simulation. At intervals of 100 simulation time steps they are dynamically updated, in order to attain the most favourable position, effectively “accommodating” its lobe positions with respect to its neighbours. During a vector update step, the preferred directions of extension are matched to the set of directions the current shape of cells presents the strongest level of extension.

The initial cell positions within the field were randomly chosen. Simulations were run for 10000 time steps (see an example in Movie S5). Parameters used for each used specified cell shape are given in Table S1.

## Acknowledgements

We thank Enrico Coen for stimulating discussions, and John Fozard for critical reading and helpful comments. This work has been supported by Consejo Nacional de Ciencia y Tecnologia (CONACYT) and by the UK Biological and Biotechnology Research Council (BBSRC) via grant BB/J004553/1 to the John Innes Centre. VAG acknowledges support from the Royal Society Dorothy Hodgkin fellowship. JAvR acknowledges support from the Netherlands Consortium for Systems Biology (NCSB), which is part of the Netherlands Genomics Initiative/Netherlands Organization for Scientific Research (NGI/NWO).

## Supplementary Materials and Methods

### Decomposing shape: Lobe Contribution Elliptical Fourier Analysis (LOCO-EFA)

In this section, we first summarise previous efforts to make EFA coefficients interpretable within a morphometrics perspective and explain why matching EFA coefficients with shape features generally does not hold. We describe in detail our new method, Lobe Contribution Elliptical Fourier Analysis (LOCO-EFA). We show how it provides quantitative and biologically interpretable measurements that are unique for a given shape, overcoming the shortfalls of the previous methods.

## Standard Fourier Analysis cannot be used to quantify complex cell shapes

Standard Fourier Analysis has been widely used to analyse cell morphology. It can, however, only be applied when cells present simple holomorphic shapes, i.e., when the radii emanating from the centroid of a cell intersect its outline only once (Figure S2A and Pincus and Theriot (2007)). When the geometry of a cell is more complex, as in the case of pavement cells, and radii emanating from the centroid can intersect the outline more than once, the shape cannot be decomposed using a Fourier expansion based on polar coordinates (Figure S2B and Schmittbuhl et al. (2003)).

## Elliptical Fourier Analysis fails to align mode frequency with morphological features

In 1982 Kuhl and Giardina proposed the Elliptical Fourier Analysis (EFA) to describe the contour of any two-dimensional shape (both holomorphic and non-holomorphic), derived from the coordinates of all the points along its outline.

In short, EFA takes the *x* and *y* coordinates of a closed contour and decomposes it into an infinite summation of related ellipses:

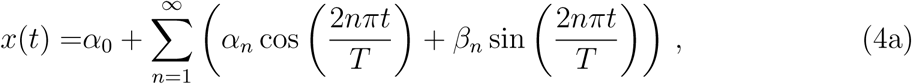

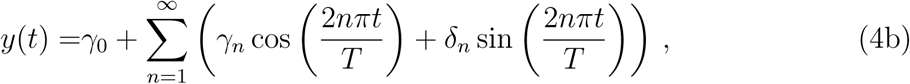

where *α*_*n*_, *β*_*n*_, *γ*_*n*_ and *δ*_*n*_ are the so-called EFA coefficients and *α*_0_ and *γ*_0_ are the *x*and *y*-offset of the initial contour. The detailed derivation of the formulae for *α*_0_, *γ*_0_, *α*_*n*_, *β*_*n*_, *γ*_*n*_ and *δ*_*n*_ can be found in Kuhl and Giardina (1982). They are calculated from a discrete chain of contour points (*x*_*i*_*, y*_*i*_) with *i* = 1*, …, K* (see Figure S3A), *K* being the total number of points along the closed contour. We define (*x*_0_*, y*_0_) *≡* (*x*_*K*_*, y*_*K*_), given that cell contours are closed. Now imagine drawing the contour of the cell, then *Δt*_*i*_ is the time spent drawing the line segment of the contour that links (*x*_*i-*1_*, y*_*i-*1_) to (*x*_*i*_*, y*_*i*_), i.e., 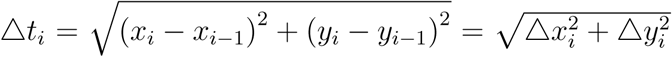. Note that *Δt*_*i*_ is not fixed but can vary for each interval. Define *T* as the total time spent to draw the whole contour, i.e., 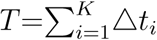. The “time” passed while drawing the contour, starting from contour point (*x*_0_*, y*_0_), or, equivalently, the distance passed along the contour to reach each contour point (*x*_*i*_*, y*_*i*_), is referred to as *t*_*i*_, i.e., *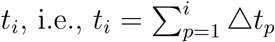*, with *t*_0_ = 0, and *t*_*K*_ = *T*, the “total drawing time” or total perimeter length (see Figure 1 in the main text). Given that no equal spacing between the points is required, it is straightforward to define *K* observation points from any kind of cell contour. The only requirements are that the contour is closed and the coordinates form an ordered list that follows the contour. The EFA coefficients are then given by:

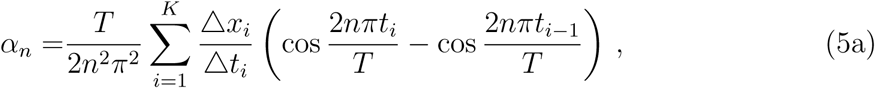

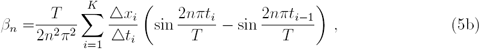

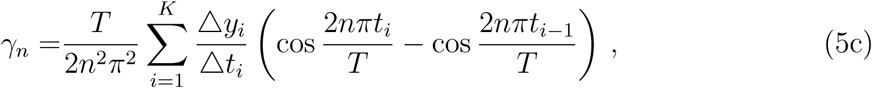

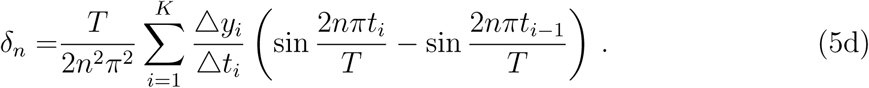

The offset to the contour is given by:

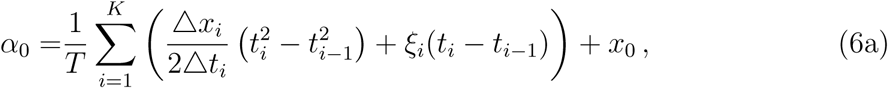

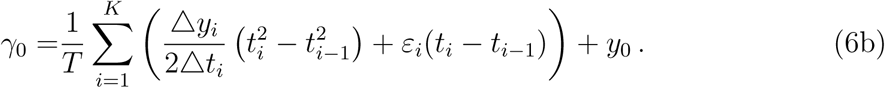

where *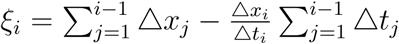* 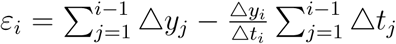, and *ξ*_1_ = *ε*_1_ = 0. Further details and full derivation can be found in Kuhl and Giardina (1982).

Each set of four coefficients yields an ellipse (also referred to as the “*n*th mode” or “*n*th elliptic harmonic”), with a certain orientation and a certain starting point. The original cell outline can thus be expressed as an infinite summation of ellipses. Note that *x*(*t*) and *y*(*t*) are periodic functions with period equal to *T*.

A visual way to understand how the set of ellipses gives rise to the final shape is as follows: the second elliptic harmonic traces two clockwise or counter-clockwise revolutions around the first harmonic; the third harmonic traces three revolutions around the path drawn by the second harmonic; and the *n*th harmonic traces *n* revolutions around the path drawn by the previous harmonic (see Figure S3 and Movie S1).

Diaz et al. (1990) proposed a heuristic measure regarding the contribution of each harmonic to the shape through an approximation of the perimeter of each ellipse multiplied by its harmonic number *n*:

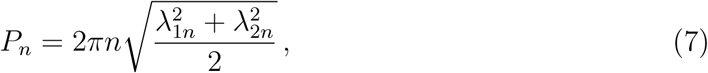

where *λ*_1*n*_ and *λ*_2*n*_ are the major and minor axis of the *n*th ellipse. Moreover, Diaz et al. (1990) introduced an additional correction to capture the complex relationship between EFA modes and shape feature periodicity.

The direction of rotation of the *n*th harmonic ellipse is given by the determinant of the EFA coefficients matrix, 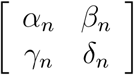, i.e., the direction of rotation is given by the sign of

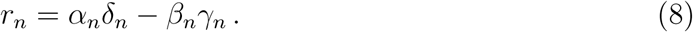

If *r*_*n*_ *<* 0, the elliptic harmonic is rotating clockwise; if *r*_*n*_ *>* 0 the elliptic harmonic is rotating counter-clockwise. When EFA is used for shape approximation, mode *n* contributes to shape features with an *n* + 1 or *n* − 1 periodicity. This is in contrast to standard Fourier Analysis, in which mode *n* contributes to shape features with an *n* periodicity. Standard Fourier Analysis, however, is only possible for holomorphic shapes, and hence cannot be applied to, for example, pavement cells. Diaz et al. (1990) observed that whether mode *n* predominantly contributes to shape features with an *n* + 1 or with an *n* − 1 periodicity strongly depends on whether the *n*th harmonic rotates together with or against the direction of the first harmonic (see Movie S2 and Movie S3). This effect of presenting contributions to the *n* + 1’th and *n* − 1’th mode depending on the rotation direction of the first and *n*th harmonic is a common phenomenon observed for objects orbiting around others (hereafter referred to as the relative direction effect). A well-known example of the relative direction effect is the rotation of the Earth and its movement around the sun. The actual number of rotations our planet makes per year (as observed from “star-rise to star-rise”, the so-called sidereal days) is one off from the number of days we perceive in a year (from “sunrise to sunrise”, the so-called solar days). Because our planet rotates around its axis in the same direction as it moves around the Sun, the number of solar days per year is 365, one less than the number of sidereal days per year, which is 366. If the rotation of Earth would have been in the opposite direction as its movement around the sun, the number of solar days per year would instead have been 367. In light of exactly the same principle, Diaz et al. (1990) introduced that when the *n*th elliptic harmonic is moving in the same direction as the first harmonic, its shape contribution *P*_*n*_ should be assigned to *n* − 1; inversely, when the direction of a given mode is opposite to the first harmonic, its shape contribution *P*_*n*_ should be assigned to *n* + 1.

We will show below that this simple heuristic is reasonable as long as the ellipse marginally deviates from a circle, but is is not valid in general. When the aspect ratio of the ellipse (*λ*_1*n*_*/λ*_2*n*_) is large (i.e., the elliptical harmonic is very flat, deviating significantly from a circular shape), the proposed rule fails to apply. Figure S3F illustrates a situation when the rotation direction of the first and third harmonic are opposite (and no other modes are used), yet instead of generating a contour with *n* − 1 = 2 protrusions, as expected from the heuristic rule, a four-sided outline is generated, clearly illustrating that this method of *P*_*n*_ shifting does not work in general (see also Movie S4). Moreover, it is not possible to reconstruct the original shape using the *P*_*n*_ values, and therefore cannot be used for additional analysis based on shape reconstruction as presented in the main paper. This strongly limits usage of EFA for biological shape interpretations and statistical population analysis. Surprisingly, although EFA has been used to quantify morphology at the organ level, relative direction effect has typically been ignored altogether (Yoshioka et al., 2005; Frieß and Baylac, 2003; Neto et al., 2006; Iwata et al., 1998, 2010; Chitwood et al., 2013).

Realising that the source of the problem is linked to the eccentricity of the ellipses, it became clear to us that we could overcome this issue by essentially decomposing each ellipse into two counter-rotating circles. All circles can then be redistributed, forming a new base. The details of how to do so are discussed below. We call the new base *ℒ*_*n*_, which, when summed up, can also reconstitute the original shape.

**Contouring the limitations: Lobe Contribution Elliptical Fourier Analysis (LOCO-EFA)**

To capture the biologically relevant cell shape features, overcoming the limitations of using *P*_*n*_ and rotation-dependent *n* + 1, *n* − 1 adjustments, we have developed an alternative method coined Lobe Contribution Elliptical Fourier Analysis (LOCO-EFA). As the name indicates, it correctly maps the contribution of each mode/harmonic to the corresponding morphological features. This is done by separating each elliptic harmonic into two circular harmonics, each rotating in an opposite direction.

First we rewrite the EFA (Equation 4) in matrix form:

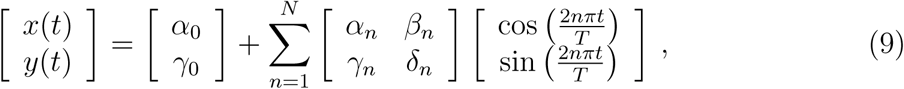

with the infinite sum being truncated at the *N* th order harmonic.

Equation 9 can concisely be expressed as

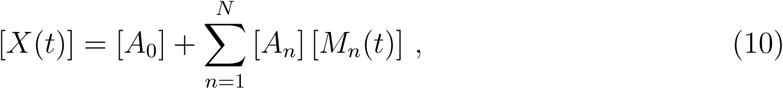

in which [*X*(*t*)] corresponds to the drawn cell outline 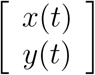; [*A*_0_] represents the spatial offset 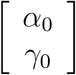; [*A*_*n*_] corresponds to the EFA coefficients matrix 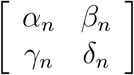; and [*M*_*n*_(*t*)] refers to the rotor 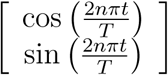. (For clarity, we will use the notation [..] throughout to emphasise we are dealing with matrices; not to be confused with *|..|* that represents determinant, which we here only refer to as det [..].)

The LOCO-EFA method consists of three steps: 1) eliminate multiple representations of the same outline; 2) decompose each *nth* elliptic harmonic into two circular harmonics, each rotating in an opposite direction; and 3) determine *ℒ*_*n*_ and *L*_*n*_ for all *N* modes. Below we describe these steps in detail.

**(1) Eliminate multiple representations of the same outline**

It had already been noted that EFA coefficients are redundant and therefore compromise statistical analysis and shape comparisons (Haines and Crampton, 2000). We found that there are three sources of degeneracy in the EFA coefficients that therefore have to be eliminated. First, a contour can be drawn starting from any arbitrary initial point along the contour. While exactly the same outline is drawn, each starting point is represented by a completely different set of EFA coefficients for all modes [*A*_*n*_]. Basically, whenever the starting point is changed, all elliptic harmonics take a different orientation (Kuhl and Giardina, 1982). The first step is therefore to transform the EFA coefficients such that the starting point of the first harmonic is always positioned at, for standardisation, the extreme of the semi-major axis (see further below). The second source of degeneracy, however, is that such a normalisation still allows for two possible representations of the outline, since each of the two extremes along the semi-major axis can be chosen as the starting point. Moreover, a third source of degeneracy is due to the fact that the outline can be drawn clockwise or counter-clockwise. Clearly, all three sources of degeneracy have to be removed to make any comparison between cells sensible.

The first step is to determine where the new starting point should be positioned, as well as the scaled amount of time or temporal angle 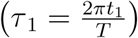 required to reach the starting point (see Kuhl and Giardina, 1982). As stated above, we wish the starting point to coincide with one of the extremes of the semi-major axis of the first harmonic. Points along the first harmonic (*x*_1_*, y*_1_) can be described as :

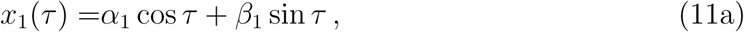

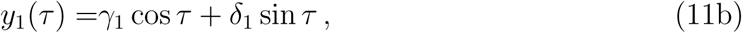

With 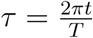 being the scaled time or temporal angle. By differentiating the magnitude of the first harmonic ellipse *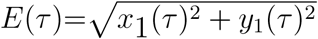* and setting its derivative to zero 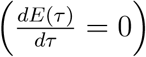, the temporal angles can be found at which the extremes along the semi-major and semi-minor axes of the first harmonic are reached (Kuhl and Giardina, 1982):

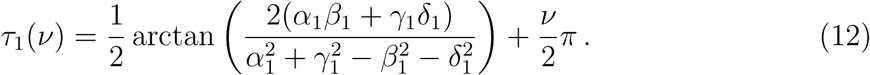

The values *v* = 0, 1, 2, 3 give the four possible solutions along both the axes, after which the same points get repeated. For LOCO-EFA, it is required (see further below) to limit the starting point to the semi-major axis only. To satisfy this condition, the second derivative of *E*(*τ*), evaluated at the temporal angle, should be negative, i.e., 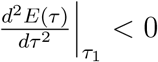. Substituting the found solutions into the second derivative results in *v* = 0 and *v* = 2 belonging to the points along the semi-major axis whenever the denominator of the arctan term is positive, and the solutions *v* = 1 and *v* = 3 belonging to the points along the semi-major axis whenever the denominator of the arctan term is negative. A very straightforward computational implementation of this result is to make use of the four-quadrant inverse tangent function (atan2) as provided by most programming languages (i.e., such that atan2(1, 1) = *π/*4 is different from atan2(−1, −1) = −3*π/*4). Then, using

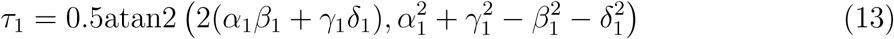

automatically and unambiguously ensures that the temporal angle *τ*_1_ is located at one of the extremes of the semi-major axis.

This still leaves two ways to position the starting point (one for each of the extremes of the semi-major axis) and thereby two distinct representations of a same outline. We therefore further restrict *τ*_1_ to always lie within the first or second quadrant (I or II in Figure S9A). This is achieved by testing if the obtained *τ*_1_ that shifts the starting point of the first harmonic to the semi-major axis indeed positions it within quadrant I or II. To shift the starting point we first introduce a time shift *τ′* = *τ -τ*_1_ such that at *τ′* = 0 the first harmonic is positioned along its semi-major axis:

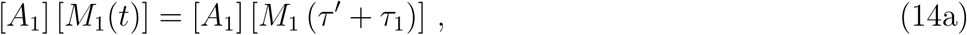

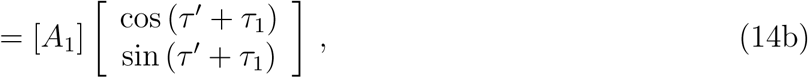

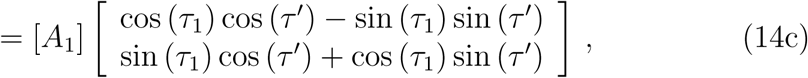

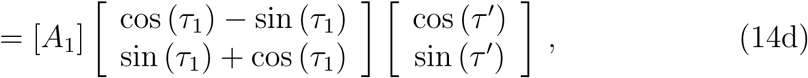

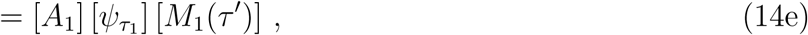

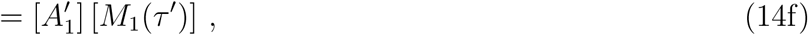

were [ψ_*τ1*_] is the rotation operator, rotating by an angle *τ*_1_ and 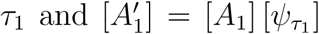. The spatial angle at the shifted starting point *ϱ* is given by

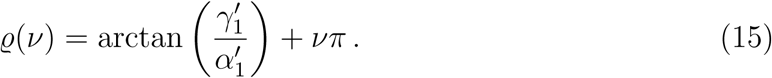

Again, a single, unique and correct solution for *ϱ* can be obtained by using *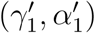* instead. The starting point lies in quadrant III or IV when *ϱ <* 0. In that case, *τ*_1_ is modified as follows:

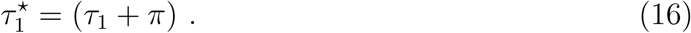

Otherwise (when the starting point is already in quadrant I or II), *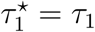*.

The new EFA coefficients corrected for the starting point then become:

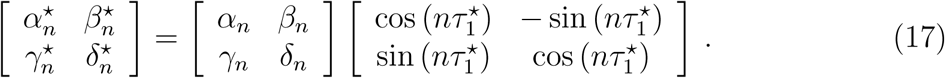

Finally, we ensure that the direction of contour approximation of the first harmonic is always counter-clockwise (i.e., that *r*_1_ ≥ 0, Equation 8). Besides removing redundancy by restricting the freedom of choice regarding the overall direction of contour approximation, this transformation also guarantees a unique correspondence between the properties of each subsequent harmonic and its contribution to the morphological features. When the direction of the first harmonic is clockwise (*r*_1_ < 0), we therefore invert the direction of motion of all ellipses, maintaining thereby their inter-relationships. This can be done by running “time” backwards:

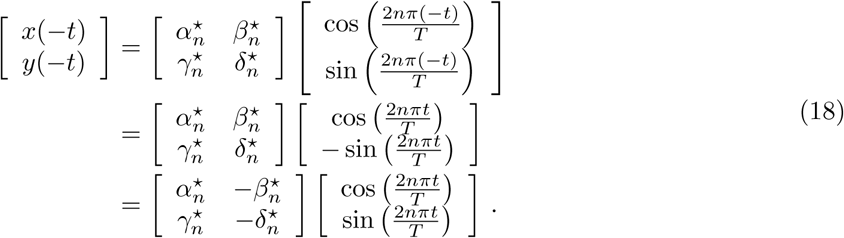

In short, whenever *r*_1_ < 0, all indices 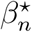 and *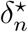* should be negated. After these steps, each unique cell contour is represented by a unique set of EFA coefficients. Note that the steps above do not alter the layout, nor do they rotate the shape. In certain study contexts, however, it might be desirable to rotate the contour itself, positioning the semi-major axis, for example, to be parallel to the x-axis (or in any other preferred orientation). The details on how to perform those rotations can be found in Kuhl and Giardina (1982). Please note that unlike in their study, our subsequent analysis does not require such a cell contour realignment.

For simplicity of notation in the rest of the Supplementary Materials and Methods we refer to the [*A*_*n*_] matrix, which elements have been normalised regarding the starting point and direction of reconstruction of the first harmonic:

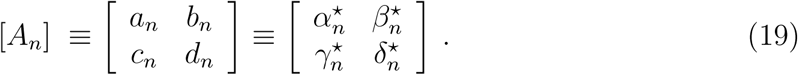

After all possible sources of redundancy have been removed, the next step of the LOCO-EFA method is to split each elliptic harmonic into two counter-rotating circles.

**(2) Decompose each** *n***th elliptic harmonic into two circles with opposite direction of rotation**

In order to find the contribution of *n*th harmonic to a given morphological feature, we rewrite the [*A*_*n*_] matrices in Equation 9 such as to explicitly introduce the length of the semi-major and semi-minor axis of the *n*th ellipse (*λ*_1*n*_ and *λ*_2*n*_).

For this purpose, it is necessary to introduce both the temporal and the spatial rotation operator of each elliptic harmonic, given by *φT*_*n*_ and *φS*_*n*_, respectively. The temporal operator is defined as

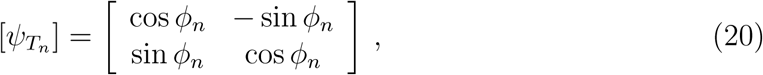

and the spatial operator is defined as

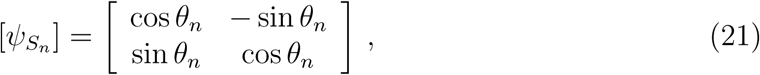

where *φ*_*n*_ is the temporal angle (i.e., the time *τ*_*n*_ required to rotate to the semi-major axis) and *θ*_*n*_ the spatial angle (i.e., the angle of this position along the semi-major axis with the positive x-axis) (Figure S9A).

Equation 10 can be written as:

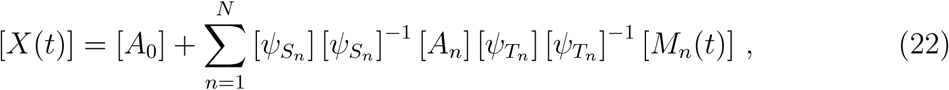

given that [*θS*_*n*_][*θS*_*n*_]^−1^ and [*θT*_*n*_][*θT*_*n*_]^−1^ correspond to the identity matrix [*I*].

Equation 10 can also be written in a form which directly highlights the contribution of the semi-major and semi-minor axis:

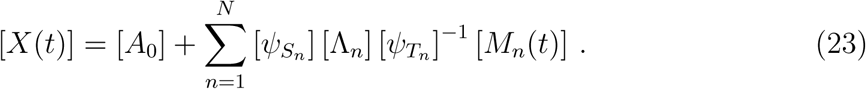

This equation can be understood as follows: for each mode, correctly position the starting point relative to the semi-major axis, transform the original circle into an ellipse, its semi-major axis along the x-axis and semi-minor axis along the y-axis, and finally rotate the ellipse to its correct position (see Figure S9B–D). Since both descriptions should be equivalent, by combining Equation 22 and Equation 23, the [λ_*n*_] matrix can be identified, with *λ*_1*n*_ corresponding to the length of the semi-major axis, and the modulus of *λ*_2*n*_ to the length of the semi-minor axis of the *n*th ellipse:

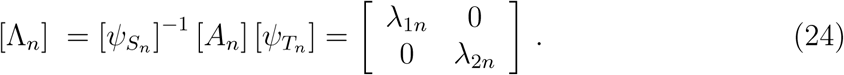

Note that this process is similar to a singular value decomposition of [*A*_*n*_], with the difference that here *λ*_2*n*_ (but not *λ*_1*n*_) can be negative. The temporal angle *φ*_*n*_ corresponds, for each elliptic harmonic, to the scaled time *τ*_*n*_ to reach an extreme along the semi-major axis (Figure S9). Similarly to Equation 12, this corresponds to

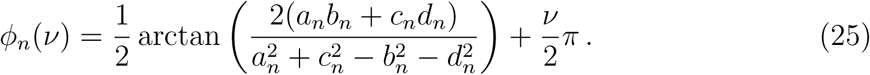

Once again, using atan2 ensures that *φ*_*n*_ corresponds to the semi-major axis (see above). For the next steps of the analysis (see below) it is essential that *φ*_*n*_ corresponds to the temporal angle to reach the semi-major axis, not the semi-minor axis, hence usage of atan2 or any equivalent function which determines the quadrant of the return value is essential.

The spatial angle *θ*_*n*_ (see Figure S9) can be calculated after applying the temporal modification:

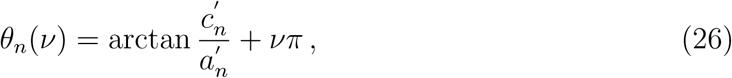

where 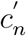 and *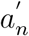*, are the new coefficients after the temporal transformation (i.e., after applying [*A*_*n*_] [*ψ*_*T*_*n*]). Again, it is essential to use atan2 to ensure that *φ*_*n*_ and *θ*_*n*_ are both relative to the same extreme of the semi-major axis. It does not matter, however, which of the two extremes is being used, hence for this step no check is required regarding the quadrants. Applying Equation 24 then provides *λ*_1*n*_ and *λ*_2*n*_. Deriving a temporal and spatial angle relative to the semi-major axis and both being related to the same extreme guarantees that *λ*_1*n*_ *≥* 0 (while *λ*_2*n*_ can be positive or negative, depending on the rotation direction of the rotor) and *|λ*_1*n*_ *| ≥ |λ*_2*n*_ *|*.

Using the above, the EFA (Equation 10) can be rewritten as:

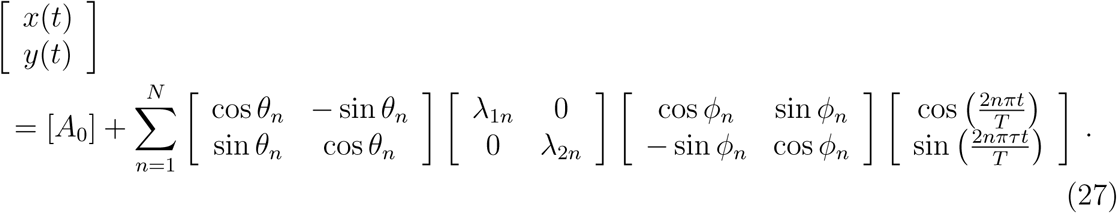

Written in this form, the contribution of each harmonic can easily be separated to correctly map to morphological feature number. The diagonal matrix containing the length of the semi-major and semi-minor axis of each *n*th mode can be decomposed into two diagonal matrices, each corresponding to circular orbits moving in opposite directions:

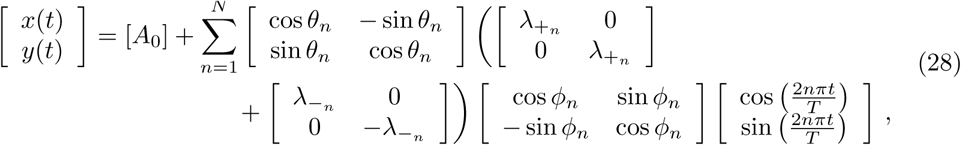

where *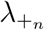*and *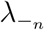* are the radii of those circles (Figure 2).

Summing up the diagonal matrices in Equation 28 yields

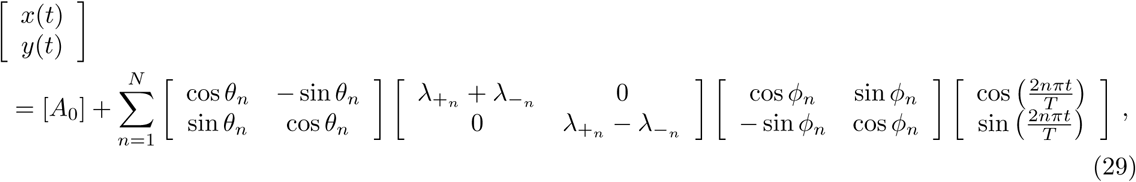

in which the major and minor axes of each elliptic harmonic are

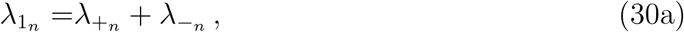

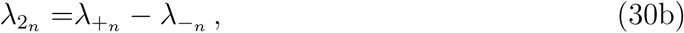

and hence the radii of each oppositely-rotating circle is given by

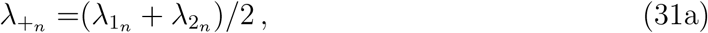

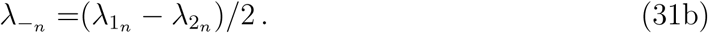

Given that *λ*_1*n*_ *≥* 0 and *|λ*_1*n*_*| ≥ |λ*_2*n*_*|*, 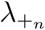 and 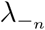 are always positive. To approximate the cell contour (*x*(*t*)*, y*(*t*)) using the circles 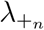 and 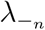 requires completing the transformations using the spatial (*θ*_*n*_) and temporal angle (*φ*_*n*_) as calculated before, most clearly seen through the expression

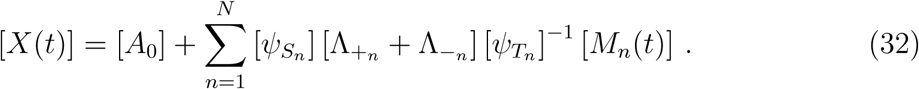

The term 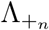 presents the subset of the *n*th elliptic harmonic which is moving in the same direction as the first harmonic, therefore purely contributing to *n* - 1 “lobes”, i.e., shape features with periodicity *n* − 1. In contrast, 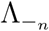 presents the subset moving in the opposite direction, purely contributing to *n* + 1 “lobes” (shape features) only. Their contributions can be separated by writing:

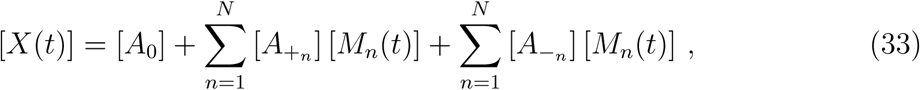

where

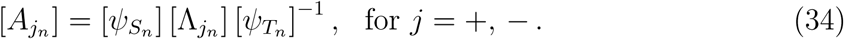

This can be further simplified. Straightforwardly, 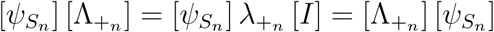. Regarding 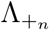,

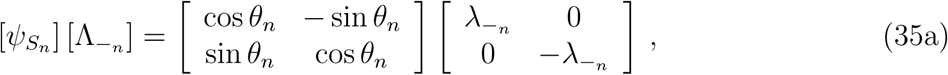

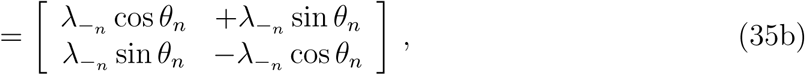

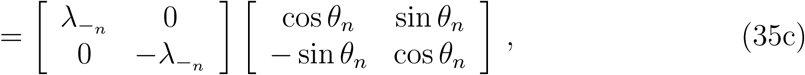

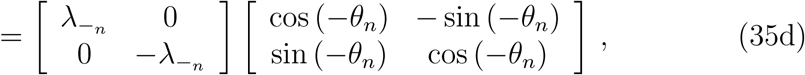

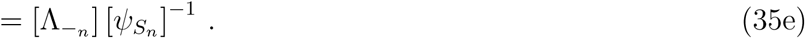

Thus,

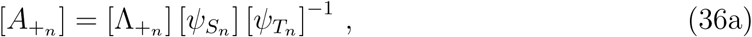

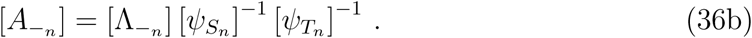

Hence, introducing *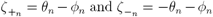*, Equation 34 can be written as

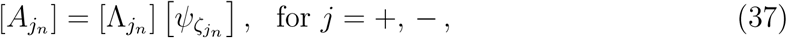

which corresponds to

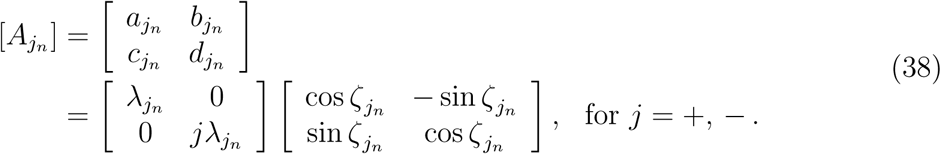

We next label those matrices with respect to their lobe contribution instead of their EFA mode. To make the distinction, we here use subscript *n* to indicate the EFA mode, and subscript *l* to indicate the LOCO-EFA mode. In general, 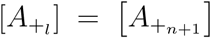 and 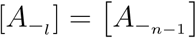. There are, however, a few exceptions, see Figure S4:

1) The overall offset of the contour is not solely given by [*A*_0_]. An additional contribution to the offset is coming from 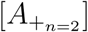. Note, however, that the contribution from 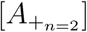 is in fact not a perfect offset to the contour, but also causes a kidney bean-shaped distortion to the contour, its deviation from a pure offset becoming more pronounced when the contribution of this mode relative to the overall contour size is larger (see Figure S4B).

2) The overall circular shape of the contour (i.e., LOCO-mode 1) receives solely a contribution from 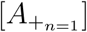 itself, i.e., 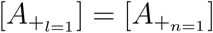.

3) When *N* EFA modes are taken into account, then LOCO-mode *N* only receives a contribution from *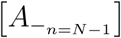*.

4) Likewise, when taking *N* EFA modes into account, there is still a contribution to LOCO-mode *L* = *N* + 1, solely coming from 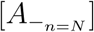.

Defining 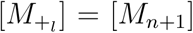 and 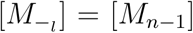, with the exceptions 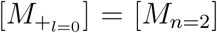 and 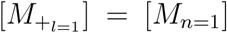, then the same shape approximation can be achieved through LOCO-EFA as through EFA:

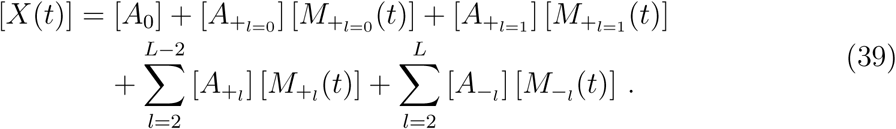

Equation 39 can be used to reconstruct the shape up to a certain *L* number, requiring EFA coefficients up to mode *L* + 1:

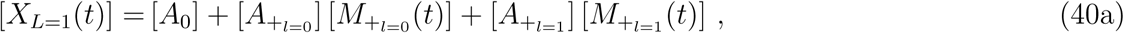

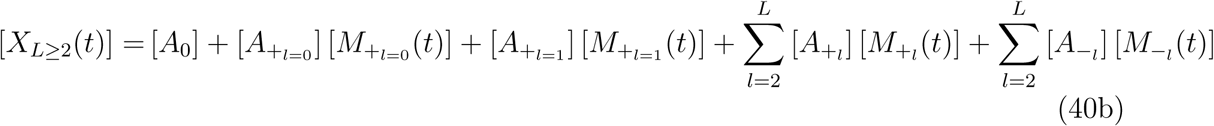

Each LOCO-EFA mode can be fully described by the combination of the radii and the starting positions of both circles. The radii are, as previously defined, *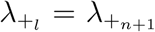* and *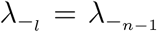*, with the exceptions *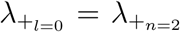* and *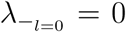*, and *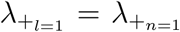* and *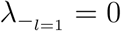*. The starting points of both circles are defined as *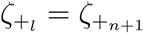* and *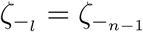*, again with the exceptions *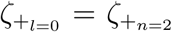* and *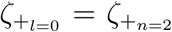*. Together this gives a set of four coefficients 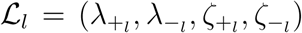 that fully capture each LOCO-EFA mode and allow for a full reconstitution of the original shape:

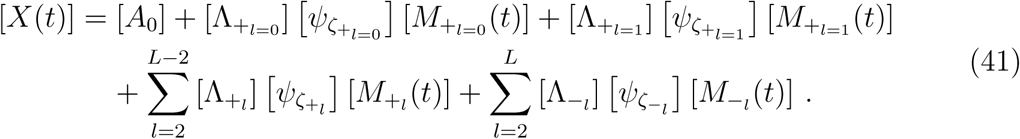

In the next section we derive how from those values a single amplitude value can be found, the *L*_*l*_ contribution.

***(3)*** *L*_*l*_ **amplitude contributions**

In this section we first show that the contribution to the shape of the two counterrotating circles of a specific LOCO-EFA mode is not simply determined by the radii of those circles, but also depends on the relative position of the starting points. We then derive a heuristic which optimally captures its contribution, determining as well the limits of this approximation. To illustrate how a difference between the starting points *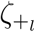* and *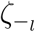* of mode *l* can affect the generated amplitude, we first plot the contribution of *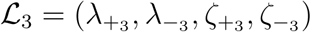* superimposed on *ℒ*_1_ = (1, 0, 0, 0), i.e., superimposed on the unit circle starting at zero degrees. As long as *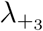*, 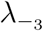 are not too large, a single amplitude value *a* (which is the amplitude of deviation from the unit circle) can be observed for any phase *ω*; peak amplitude values *A* occur for specific phases Ω (as illustrated in Figure S10I). Figure S10A–C depict the shape contribution of the negative rotor (green), positive rotor (red) and their summed contribution (orange), as a function of the phase *ω*, for _3_ = (0.15, 0.15, 0, 0). This scenario illustrates that even though the negative rotor makes two sidereal rotations while the positive rotor makes four sidereal rotations, the amplitude pattern that they generate not only has a period three in both cases, as argued throughout the paper, but also perfectly match one another regarding the phases at which the peaks and troughs in the amplitude pattern are reached. The summed contribution (Figure S10C) therefore indeed yields a peak amplitude exactly equal to *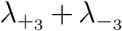*. In contrast, when both rotors are exactly out-of-phase (ℒ_3_ = (0.15, 0.15, 0*, π*), Figure S10D–F), the peaks and troughs are exactly out-of-phase as well, and the patterns almost (but not totally) cancel each other out (Figure S10F). To determine the effective contribution of a LOCO-EFA mode we therefore have to determine the phase at which each rotor reaches its peak amplitude.

The angles Ω_+_, Ω_-_ at which the positive and negative rotor reach their peak amplitude can be calculated straightforwardly. They occur when the phase of the rotor itself is equivalent to the overall phase generated by *ℒ*_1_ (as illustrated in Figure S10G). The phase of *ℒ*_1_ starts at *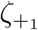*, while the phase of the positive rotor starts at *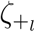* (illustrated in Figure S10K). When the pattern is laid down, the phase of the positive rotor changes (*l* + 1) times faster than the phase of *ℒ*_1_ (Equation 41). Thus, regarding the phase at peak amplitude it holds that

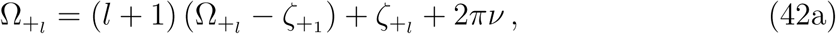

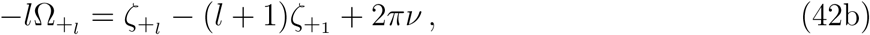

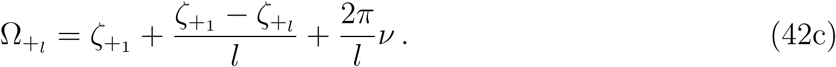

where *v* is any integer, and the values *v* = 0, 1*, …, l* - 1 provide the complete set of phases at which peak amplitude is reached. Likewise, the negative rotor rotates (*l* −1) times faster than *ℒ*_1_ and in the opposite direction, hence starting at *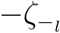* (see Figure S10J). Regarding the phase at peak amplitude it therefore holds that

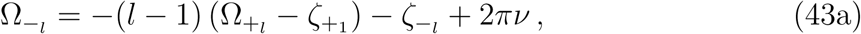

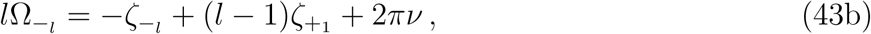

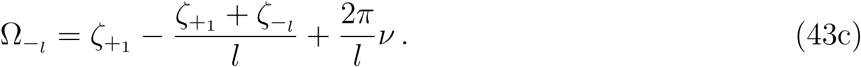

To assess the summed amplitude contribution we next fit the amplitude pattern laid down by each rotor to a sine wave, exactly matching both the peak amplitude and the phase at which the peak amplitude is reached, and then sum those two contributions:

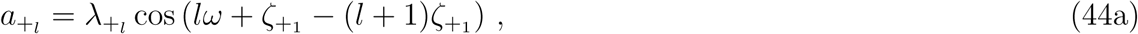

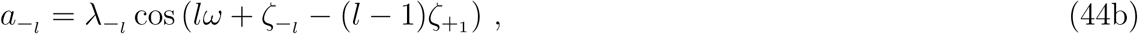

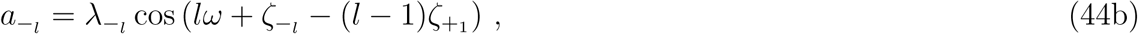

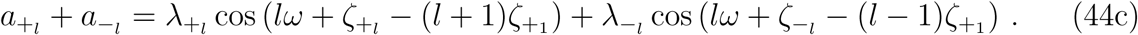

Equation 44a exactly matches the peak amplitude and its phase of the positive rotor, as derived in Equation 42, while Equation 44b exactly matches the peak amplitude and its phase of the negative rotor, as derived in Equation 43. While peak amplitude and phase match perfectly, Figure S10G, H illustrate that for not too large values of *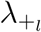*, 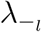 also the rest of the pattern presents a close match.

Using standard trigonometry, the summed amplitude can be written as

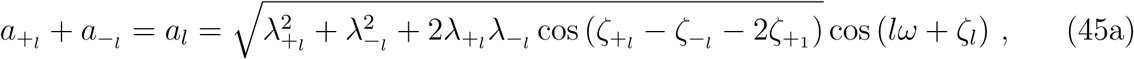

where

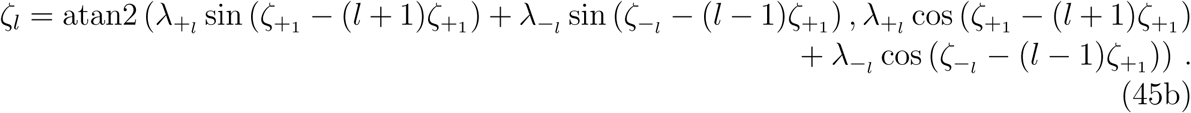

Figure S10I illustrates that for not too large values of *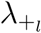*, 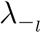 this expression provides a close match to the pattern generated by the *ℒ*_*l*_ mode, here illustrated using both a different phase and a different amplitude for both rotors. Using the equation above, the amplitude or contribution of mode *l* can hence be defined as

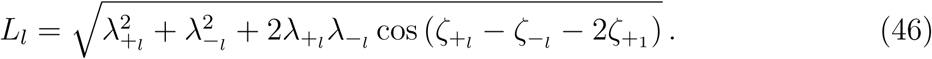

While above we have performed a more formal derivation, a more intuitive way to understand the amplitude contribution of the positive and negative rotor combined, is in terms of how much the amplitude contributions of both rotors are out-of-phase with each other. The most straightforward moment to determine their phase shift is at the initial point of drawing the cell’s outline. As illustrated in Figure S10J, K, the phase shift regarding the amplitude contribution of the negative rotor is given by *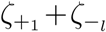*, and of the positive rotor by *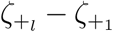*. To understand the graphical argument for the negative rotor case, one has to realise that the negative rotor rotates clockwise, and hence its starting angle is *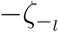* (for the positive rotor it is simply *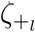*), while for the same reason the phase shift between initial amplitude and maximum amplitude has to be calculated clockwise (instead of standard counterclockwise, as done for the positive rotor), again as shown in Figure S10J, K. Plotting the phase shift and strength of both amplitude contributions as vectors and then adding them up then provides the above equation for *L*_*l*_ (as is depicted in Figure 2C).

Note that the pattern can never be perfectly described by a sine wave, which is why four parameters are needed to describe each mode, rather than the two required by Equation 45. Even when following Equation 45 the two waves cancel each other out, this is not exactly the case, as seen in Figure S10F. There is therefore no additional level of redundancy, as the usage of the *L*_*l*_ numbers might suggest, even when the value of *L*_*l*_ = 0. When the amplitude of a rotor becomes larger, the generated pattern starts to deviate from a sine wave (Figure S10J, K), and their summed pattern from Equation 45 (Figure S10L). For simple closed contours (which cell outlines should always be), the deviations cannot be very large, since otherwise the contour becomes non-simple (i.e., crosses itself). This confinement in deviations for cell outlines allows us to base a significant part of our analysis on *L*_*l*_ values.

Moreover, we observe that the positive rotor generates protrusions that are flatter than a sine wave (Figure S10B, J), whereas the negative rotor generates lobes that are more pointy (Figure S10A, K). The extent of “flatness” or “pointiness” of each mode can be determined by the proportion *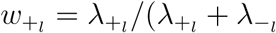* and *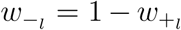*, respectively; a useful additional measure if needed.

## Entropy measurement

The entropy measure is defined as

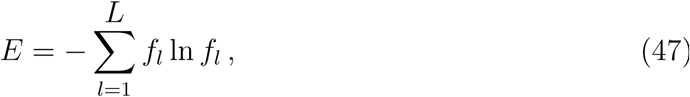

where *f*_*l*_ refers to the relative proportion of each *L*_*l*_ for a given *L* number of modes analysed, i.e., *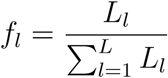*

## Final remarks

To summarise, the LOCO-EFA method consists in: 1) eliminating degeneracies in the EFA coefficients; 2) decomposing each elliptic harmonic into two circles rotating in opposite directions (*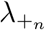* and *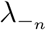*) and therefore contributing to *n* −1 and *n* + 1 number of lobes (more generally, morphological features); and 3) calculating the offset between starting points of these two circles derived from each ellipse to estimate the amplitude of the *L*_*l*_th lobe contribution.

To eliminate cell area effects (for example, when looking at shape diversity within a cell populations), it might also be desirable to normalise for cell size by dividing each *L*_*n*_ value by the square root of the cell area. This then provides a complete description of the number of lobes and their amplitude, which can be used to characterise and quantify intrinsic cell shape properties, irrespective of cell area, spacing between sampling points, and rotations or inversions of the cell (Figure S11A, B). Changes in the image resolution, however can of course affect the fine-grained information retrieval, but only if the resolution becomes very low (Figure S11C, D).

## Supporting Figures

**Figure S1:**
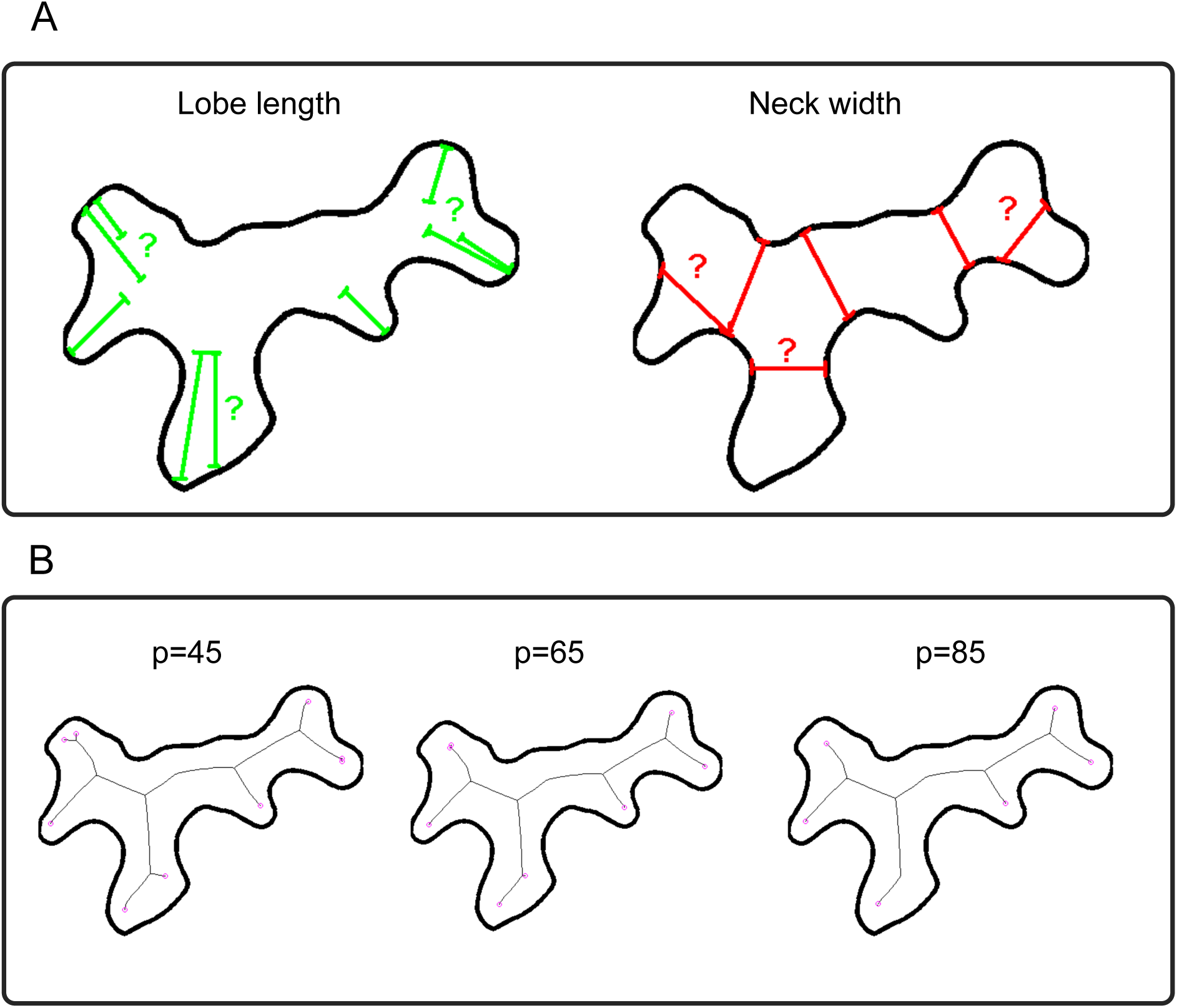
Subjective choices involved in neck width and lobe length determination and Skeletonisation. (A) Neck width and lobe length depend on human criteria for both identifying and quantifying such structures, as indicated by the question-marks. (B) Both the number of skeleton end-points and the length of the branches strongly depend on the parameter settings used for the skeletonisation algorithm. Here, parameter *p* (see material and Methods) was set to 45 (left), 65 (middle), or 85 (right). For those values, the number of predicted lobes varies between 6 and 8.

**Figure S2:**
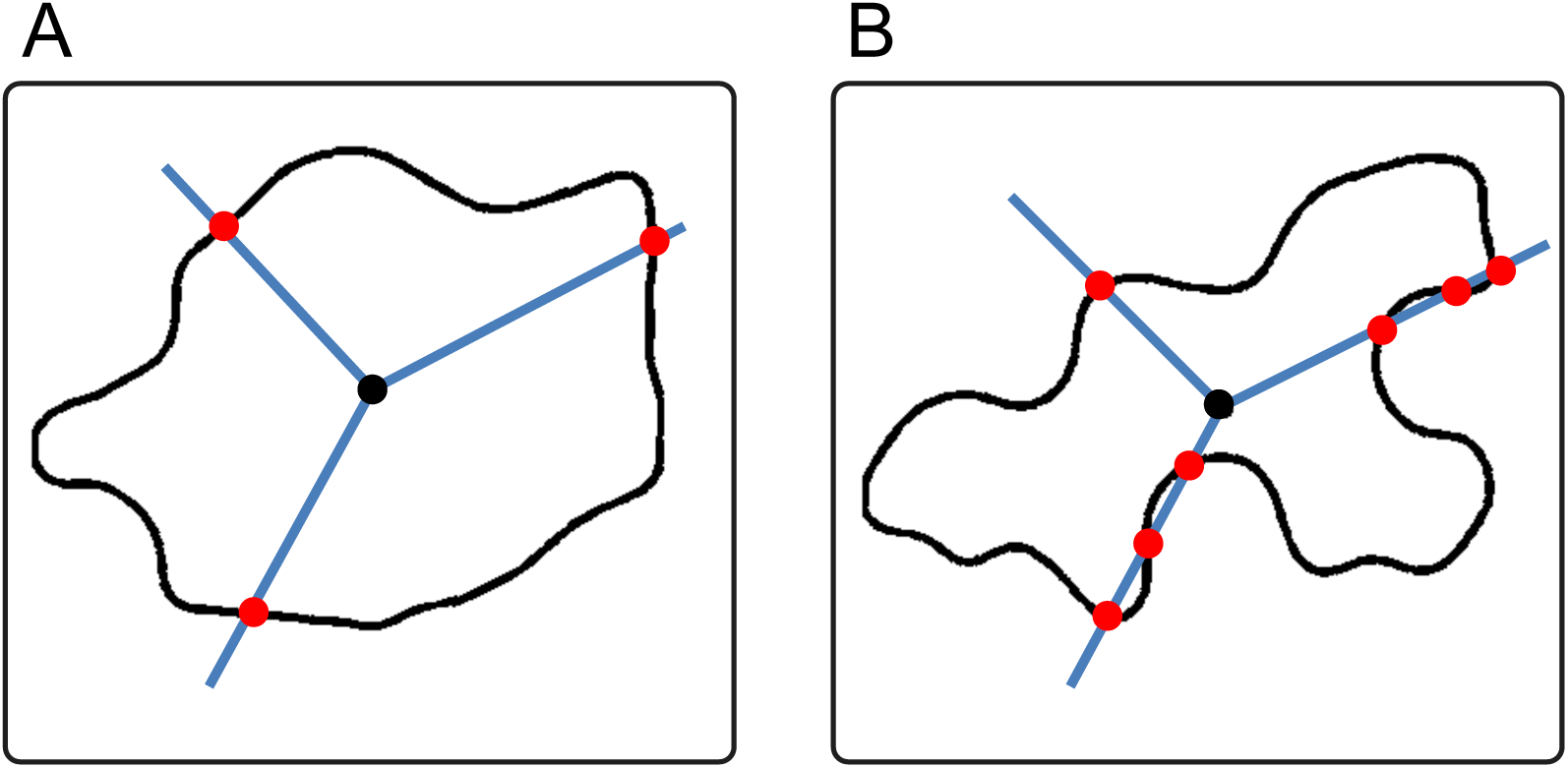
Holomorphic and non-holomorphic shapes. (A) In holomorphic shapes all radii starting from the centroid intersect the outline only once. (B) In non-holomorphic shapes some radii intersect more than once. Very few pavement cells have a holomorphic shape, the majority presenting highly complex non-holomorphic outlines. The outline of non-holomorphic shapes cannot be represented as a function of the angle, precluding, for example, standard Fourier analysis.

**Figure S3:**
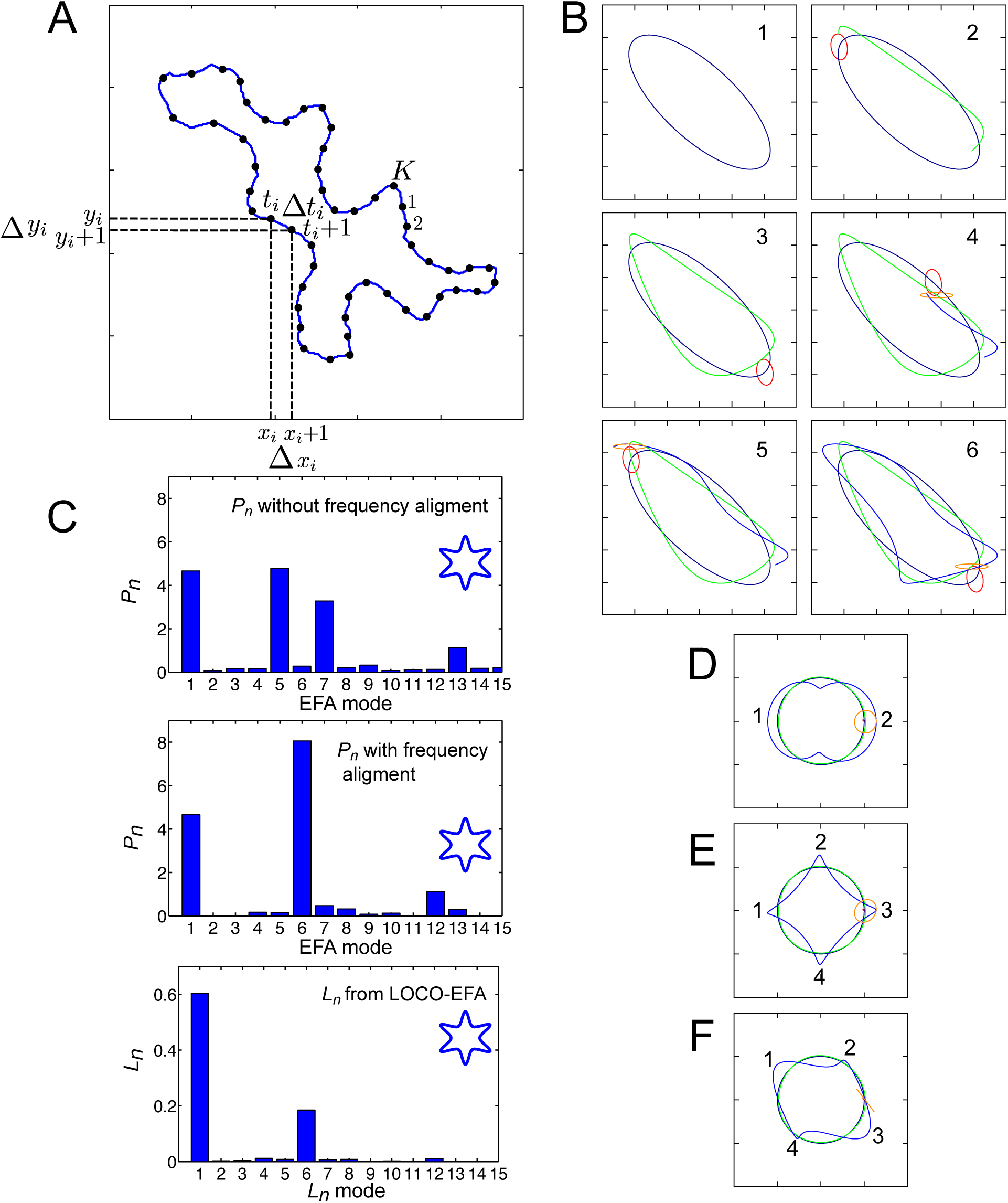
Cell outline reconstruction using EFA and LOCO-EFA. (A) Calculating the EFA coefficients from a cell outline. The discrete chain of contour points can be positioned arbitrarily (for example, the points do not have to be associated to an underlying grid). Also the distances between points can be arbitrarily long. (B) Sequential approximation of the cell’s contour. The first harmonic forms an elliptic shape (1, blue). The second harmonic describes an elliptic orbit (2–6, red), orbiting twice while moving around the first harmonic (2, 3). Their summed trajectories are shown in green. The third elliptic harmonic (4–6, orange) orbits thrice while moving around this summed trajectory (4–6). The summation of the first three elliptic harmonics is shown in blue. See Movie S1 for this dynamical reconstitution of the cell contour. (C) The EFA coefficients cannot be directly linked to shape features, here shown through the power contribution of each harmonic (*P*_*n*_) (upper panel): the main *P*_*n*_ contributions of a six-pointed star-like shape (shown as an inset) come from the 5th and 7th harmonic. To align the coefficients to actual shape features, Diaz et al. (1990) proposed to shift the contribution of each harmonic to either *n* + 1 or *n* −1, depending on the rotation direction of each individual harmonic (middle panel). This brings the main shape contributor and the actual number of shape features in alignment to each other. This method, however, does not always hold (see F), and moreover generates a range of spurious contributions from a large set of different modes, which is a notorious issue that hampers analysis of complex shapes using standard EFA (Haines and Crampton, 2000). The LOCO-EFA method correctly aligns the shape assessment and the real shape features (lower panel), without generating any additional spurious contributions. Moreover, unlike the other methods, the values correctly represent the amplitude of the shape features. (D–F) Contours (shown in blue), generated from the first (green) and third EFA harmonic (orange) only. The number of morphological protrusions (lobes) specified by the *n*th EFA mode is affected, but not fully determined, by its rotation direction. The heuristic rule proposed by Diaz et al. (1990) states that if the first harmonic and the *n*th harmonic rotate in the same direction, a contour is generated with *n* - 1 protrusions, while if their rotation direction is opposite, the contour will contain *n* + 1 lobes. This rule, however, is only correct when the elliptic harmonic has a circular shape (D, E), in which case indeed overall shapes are generated with either 2 (D) or 4 lobes (E), simply dependent on the rotation direction with respect to the first mode. When, however, the elliptic harmonic has a higher eccentricity (F), the final shape can have *n* + 1 protrusions (4 lobes) even though the rotation direction of the first and third harmonic are opposite. This is a consequence of each single EFA mode actually contributing to two different spatial modes. See also Movie S2–S4.

**Figure S4:**
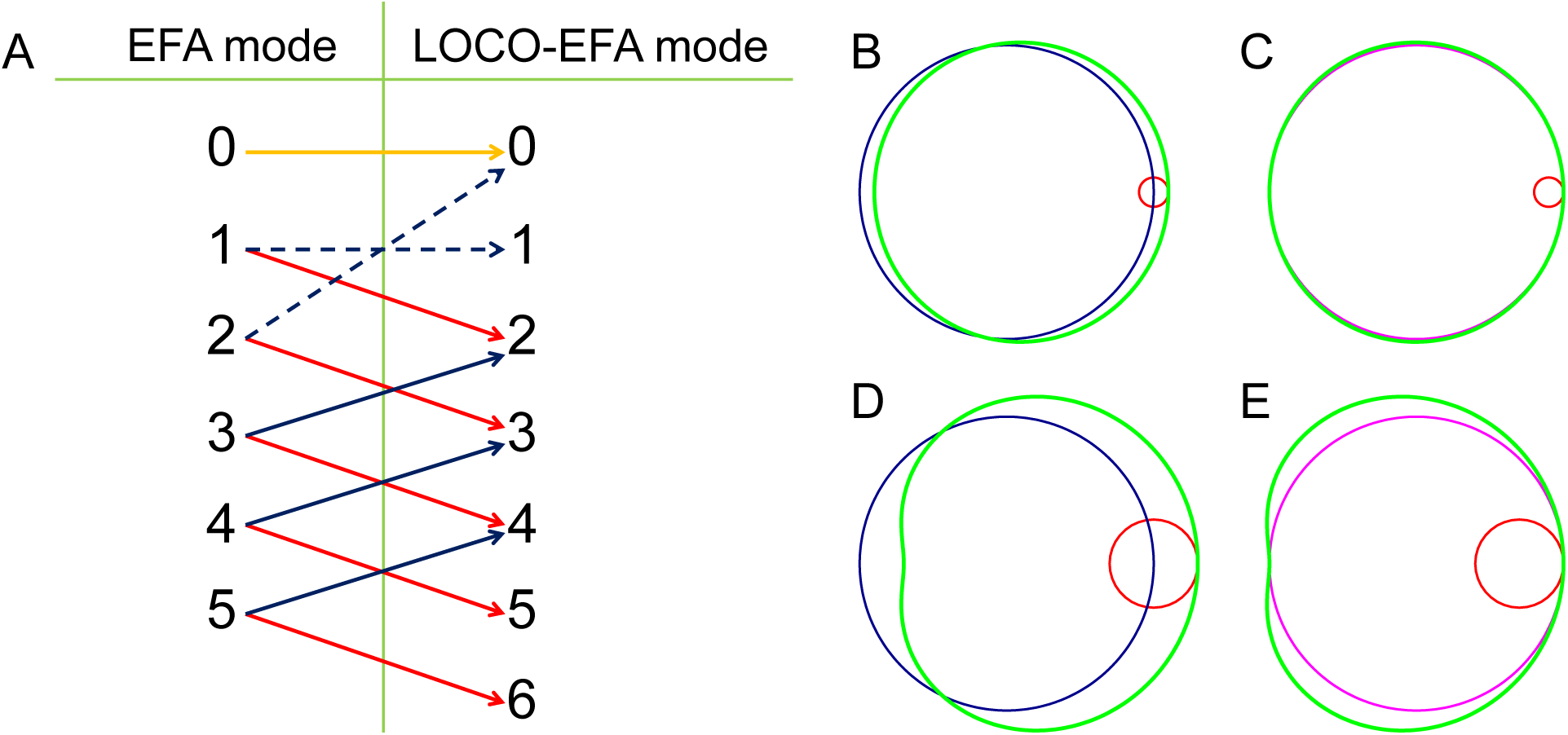
Schematic mapping between EFA modes and LOCO-EFA modes. (A)Each *n*th EFA mode contributes to both *n* + 1 and *n* −1 morphological periodicities. The red arrows represent the contributions of the *n*th EFA mode to *n* +1 protrusions, due to the clockwise rotations of the circular harmonics *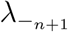*. The blue arrows indicate the contributions to *n* - 1 protrusions, due to the counter-clockwise rotations of the circular harmonics *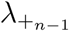*. A few exceptions apply: The second EFA mode contributes to a shift in the positioning of the layout, i.e., to LOCO-EFA mode 0 (*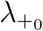*, blue and dashed line), rather than to the overall size of the layout, as might have been expected. The first EFA mode contributes to the overall circular size of the layout (*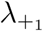*, blue and dashed line), rather than to a shift in the positioning of the layout, as might have been expected. The zeroth EFA mode only contributes to a shift in the positioning of the layout (yellow line). Finally, the two highest LOCO-EFA modes have incomplete contributions, given any cutoff in the number of EFA modes. (B–E) The contribution *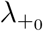* is not simply an offset of the contour, but also involves a kidney bean-shaped distortion, more pronounced for larger contributions. Mode *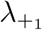* (the circular mode) is shown in blue; mode *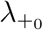* in red; their summation in green; and the predicted shape if the contribution were purely an offset in magenta. (B) *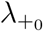* being 10% of *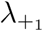*, effectively resulting in just an offset to the shape outline (C). (D, E) *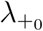* being 30% of *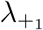*, resulting in a clear kidney bean-shaped distortion of the shape.

**Figure S5:**
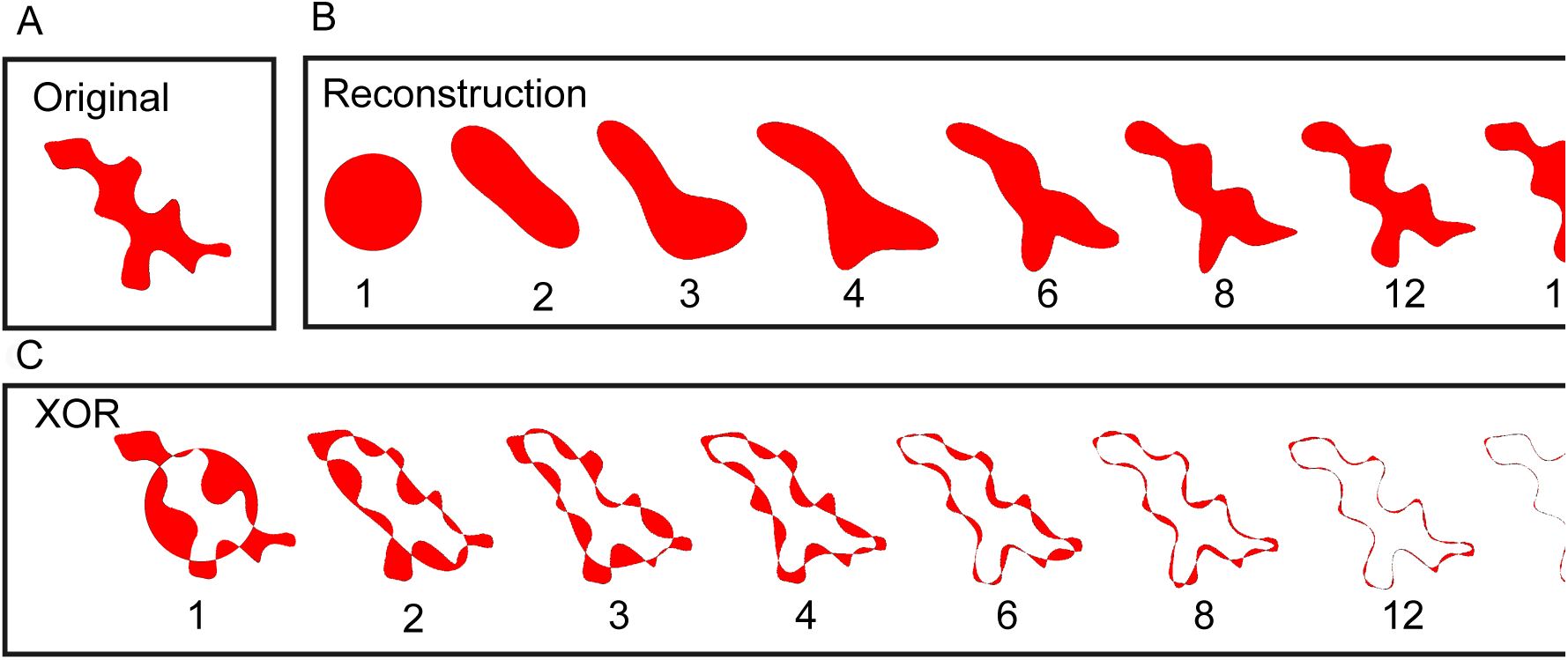
XOR measure extracted from the LOCO-EFA coefficients to provide additional information regarding the cell shape complexity. (A) Original cell contour. (B) LOCO-EFA reconstruction taking the first *n ℒ*_*n*_ modes into account, as indicated below the panels. (C) Determination of the level of mismatch between the original cell shape and its *n*th order truncated LOCO-EFA approximation, by applying an XOR (Exclusive OR) function.

**Figure S6:**
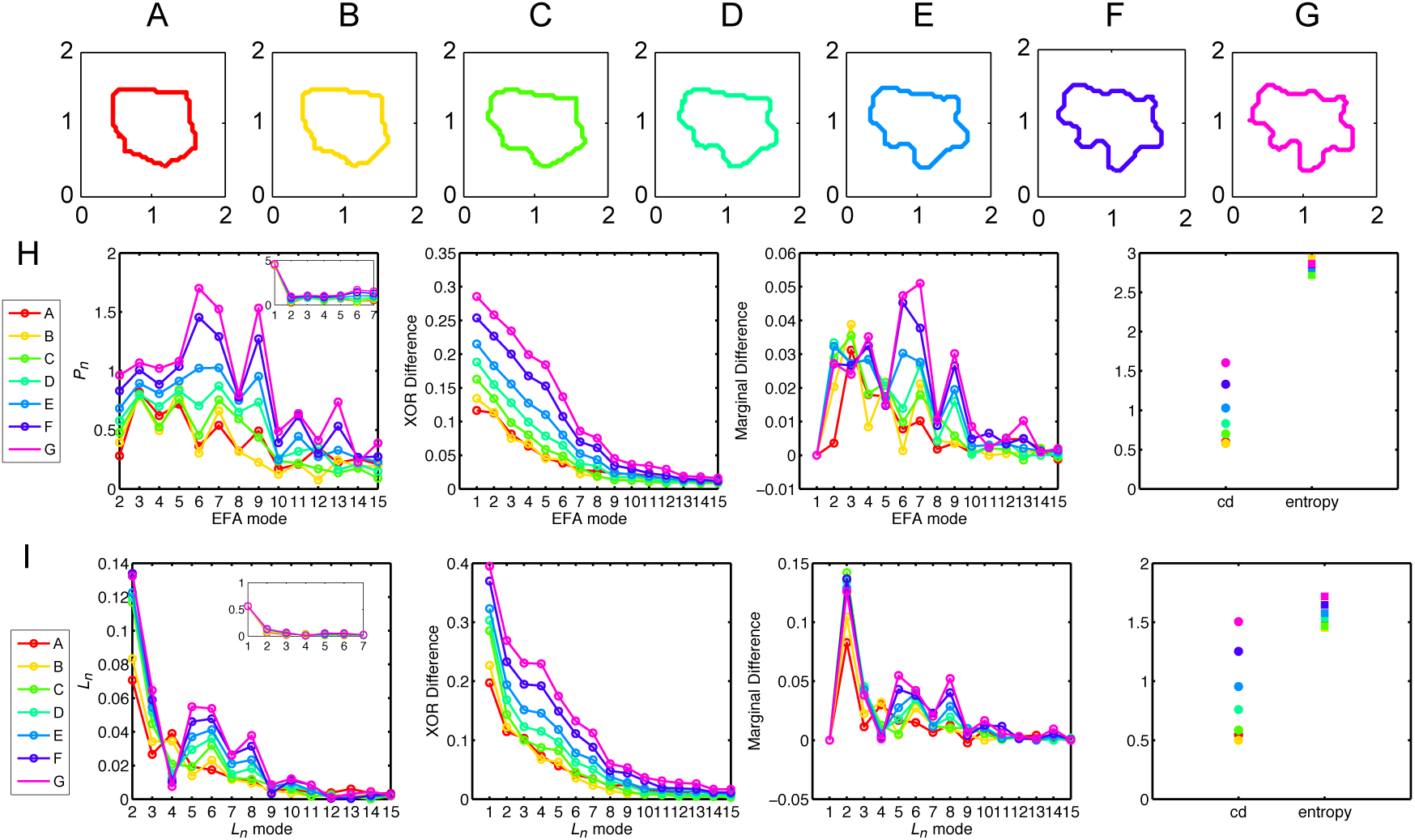
Additional example of LOCO-EFA metrics on a cell changing its shape over time. (A–G) Sequence of a tracked pavement cell growing over time with normalised area. (H) *P*_*n*_, XOR difference and marginal difference profiles, cumulative difference and entropy using EFA. (I) *L*_*n*_, XOR difference and marginal difference profiles, cumulative difference and entropy using LOCO-EFA. See also the first example presented in Figure 4 in the main text.

**Figure S7:**
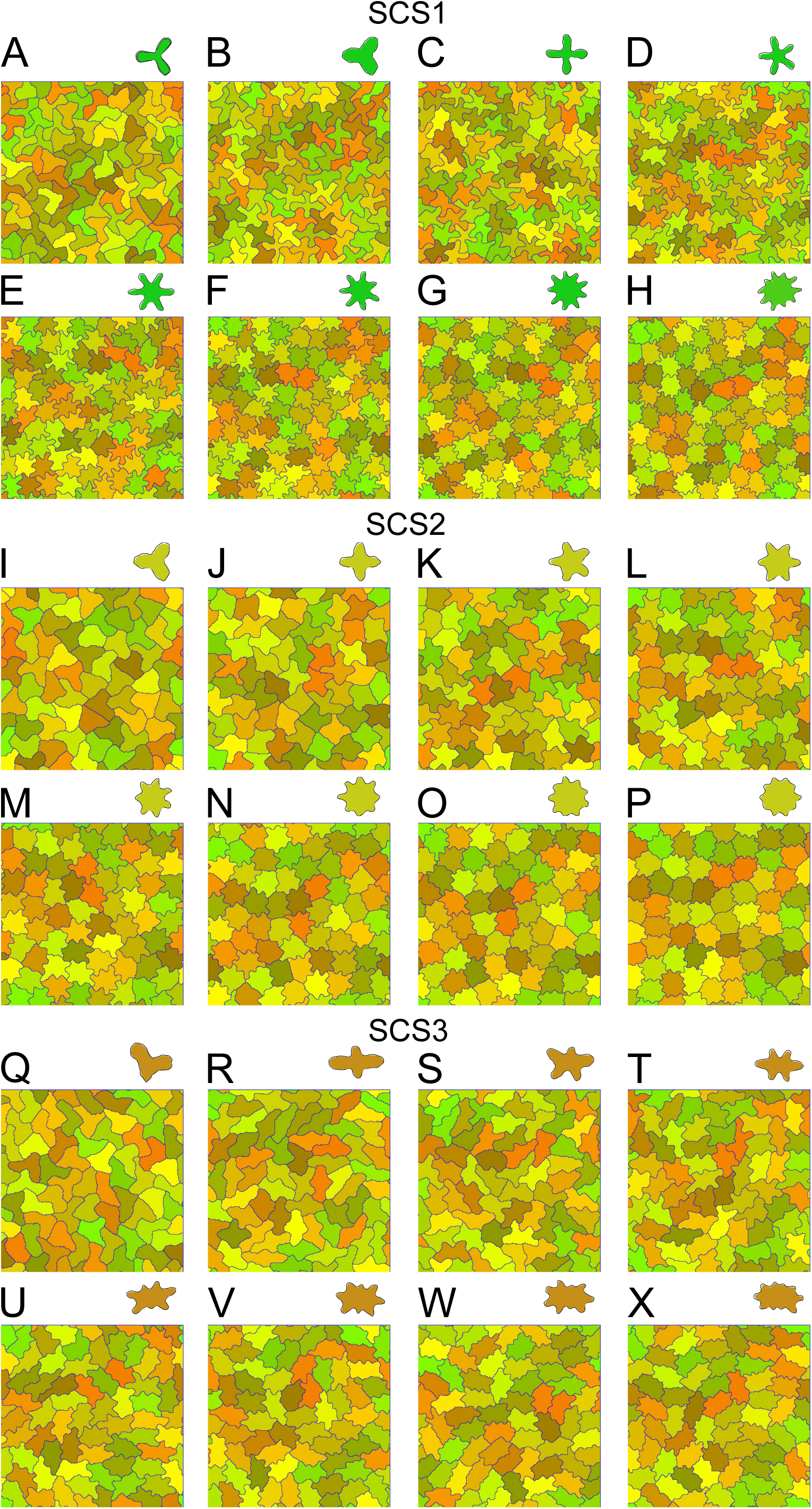
Confluent *in silico* cell populations, simulated with three types of specified cell shapes (SCSs) and different number of specified lobes. Parameters for the different specified cell shapes are given in Table S1. (A–H) Cells with large protrusions (Specified Cell Shapes 1, SCS1), with specified lobe number increasing from 3 (A) to 10 (H); (I–P) Cells with small protrusions (SCS2), with specified lobe number increasing from 3 (I) to 10 (P); (Q–X) Elongated cells (SCS3), with specified lobe number increasing from 3 (Q) to 10 (X). Specified cell shapes, as resulting from single-cell simulations, are shown above each panel. Cells are randomly coloured. See also Figure 5 and Figure S8.

**Figure S8:**
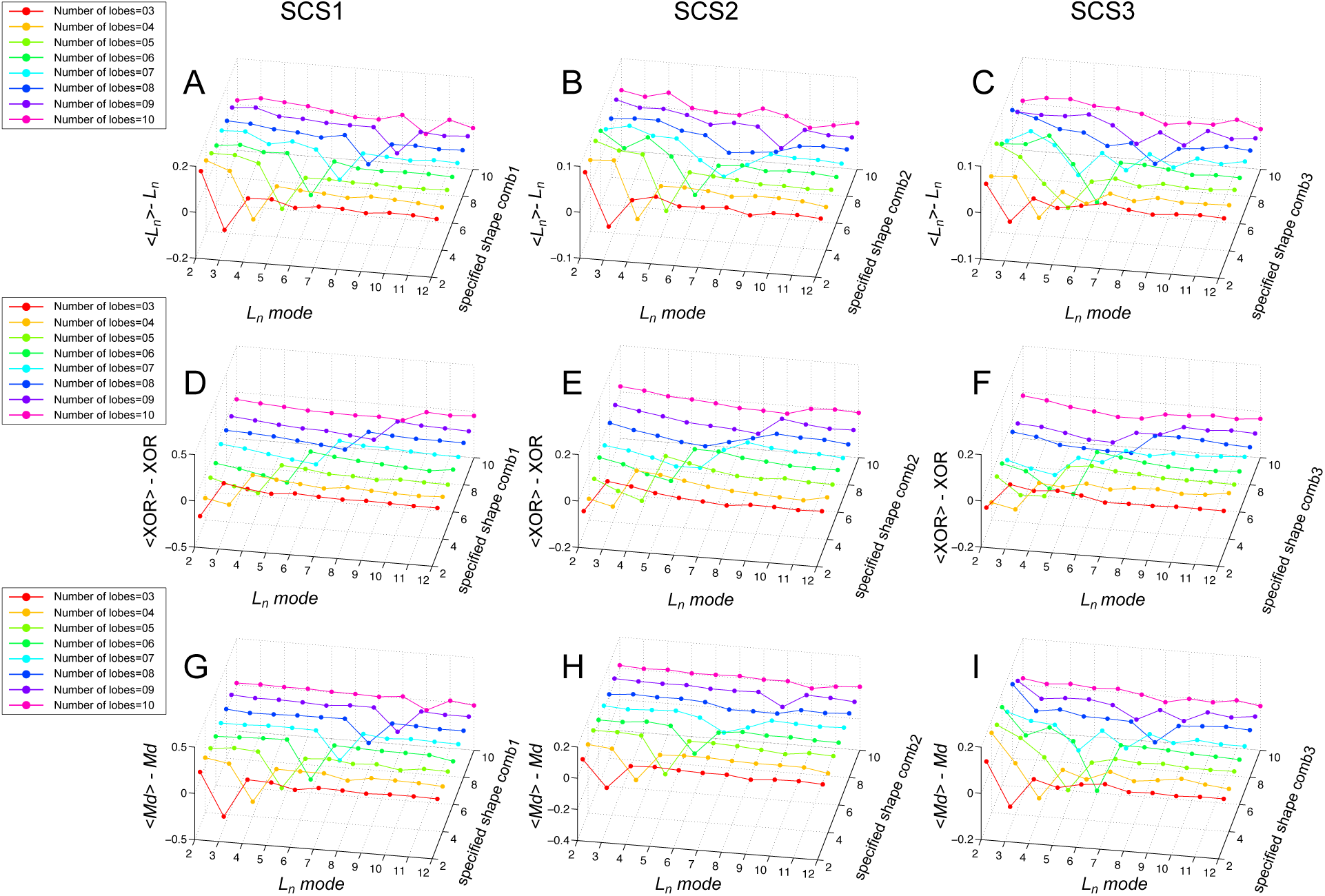
Divergence between specified cell shapes and resultant population-level cell shape diversity within confluent *in silico* cell populations. The mismatch is visualised as the difference in LOCO-EFA-derived measures between the average for confluent population simulations and its value for single cell simulations. Three types of specified cell shapes (SCSs) were simulated, as indicated for each column, and parametrised in Table S1. Lobe numbers vary from 3 to 10, as indicated for each row and depicted in Figure S7. (A–C) Difference between the average *L*_*n*_ (⟨*L*_*n*_ ⟩) values of the cells of the simulated populations and that of a single simulated cell. (D–F) Difference between average *XOR* (⟨*XOR⟩*) of the cells of the simulated population and that of a single simulated cell. (G–I) Same as in (A–C, D–F), but now for marginal difference (Md). In all cases the specified cell shape becomes less pronounced in the confluent population simulations (indicated by negative values that all measures yield at the given specified lobe number), while the cells present an increased shape diversity and complexity (seen by the broad flanking regions with positive values in the profiles, indicating a large range of modes that contribute to the shapes). These effects are present regardless of the number of specified lobes or the combination of parameters used, but becomes less pronounced at higher lobe numbers. See also Figure 5.

**Figure S9:**
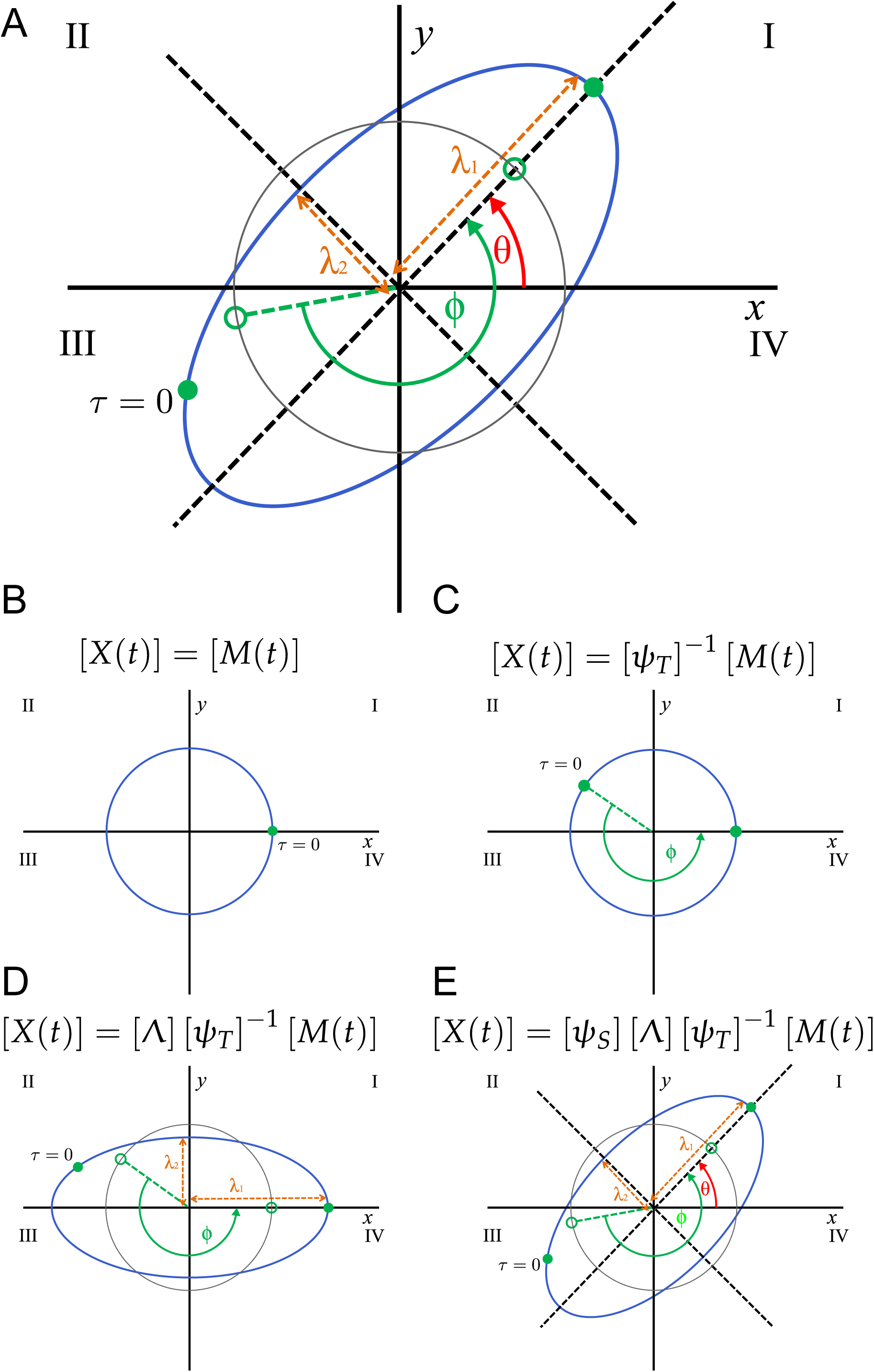
Temporal and spatial transformations required to calculate the precise contribution of each EFA mode. (A) The blue ellipse depicts an elliptic harmonic, given by [*X*(*t*)] = [*A*] [*M* (*t*)], while the grey circle depicts the unit circle, given by [*X*(*t*)] = [*M* (*t*)]. The temporal angle *φ* is the scaled time required to move along an elliptic harmonic from the starting point at *τ* = 0 to one extreme along the semi-major axis (green filled circles). The angle *φ* cannot be trivially derived from the spatial position at *τ* = 0, requiring first an effective projection upon the unit circle (green open circles). The spatial angle *θ* is the inclination of the elliptic harmonic. After applying the appropriate spatial and temporal transformations to the EFA coefficients using this geometrical interpretation, the semi-major and semi-minor axis, *λ*_1_ and *λ*_2_, can be straightforwardly obtained. To eliminate multiple representations of the same outline, the starting point of the first harmonic is specifically positioned at the extreme along the semi-major axis which lies in either quadrant I or II, given by temporal angle *τ*^*\**^ (Equation 17). In contrast, all other temporal angles *φ*_*n*_ used, while also positioned along the semi-major axis, are not confined to quadrant I or II. (B–E) Visual interpretation of Equation 23 as a step-wise construction of the elliptic harmonic. The matrix [*M* (*t*)] corresponds to the unit circle (B). For each mode, first the starting point relative to the semi-major axis is correctly positioned (C); then the original circle is transformed into an ellipse, its semi-major axis along the x-axis and semi-minor axis along the y-axis (D); and finally the ellipse is rotated to its correct position (E). See further details in the Supplementary Materials and Methods. *ℒ ℒ*

**Figure S10:**
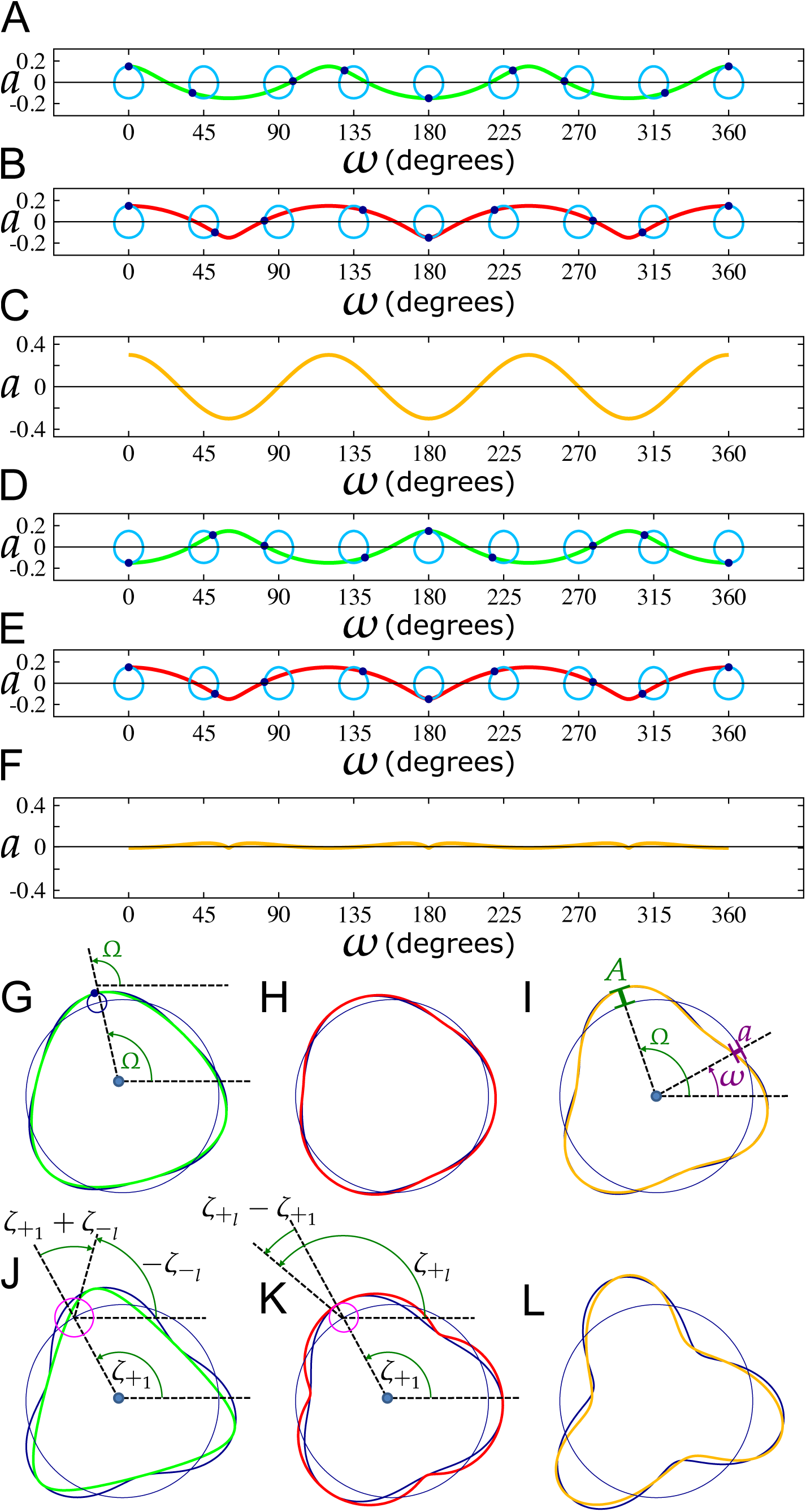
Impact of LOCO-EFA starting points *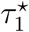* **and** *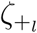* **on the amplitude of the reconstructed shape**. All panels show *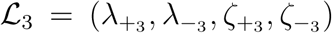* superimposed on _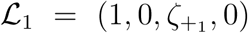_, i.e., the unit circle. In (A–F), _1_ = (1, 0, 0, 0). (A) Amplitude *a* as a function of phase *ω* (as depicted in I) for the negative rotor only (*ℒ*_3_ = (0, 0.15, 0, 0)), shown in green. For several phases also the rotor (light blue) itself is drawn. Note that by plotting the rotor in the (*ω, a*) plane, the circles are slightly deformed. (B) Same for the positive rotor only (*ℒ*_3_ = (0.15, 0, 0, 0)), in red. (C) Same for both rotors superimposed (_3_ = (0.15, 0.15, 0, 0)), in orange. (D–F) Alike (A–C), but for an out-of-phase starting angle of the negative rotor. (D) *ℒ*_3_ = (0, 0.15, 0*, π*), in green; (E) *ℒ*_3_ = (0.15, 0, 0, 0), in red; (F) *ℒ*_3_ = (0.15, 0.15, 0*, π*), in orange. In (G-L), 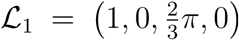 (G) Pattern generated by the negative rotor only 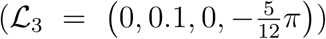, shown in green, compared to the equation *a*_-*l*_ = 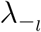 (Equation 44b), in dark blue). They present a close match. (H) Same for the positive rotor only 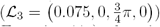, shown in red, compared to 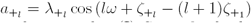 (Equation 44a), show in dark blue, again presenting a close match. (I) Same for both rotors superimposed 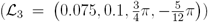, shown in orange, and *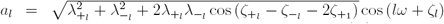,* (Equation 45), show in dark blue, again presenting a close match. Panel (G) also illustrates that at peak amplitude the phase of the main and subrotor are equal; panel (I) also illustrates the concepts amplitude (*a*); peak amplitude (*A*); phase (*ω*); and phase at peak amplitude (Ω). (J–L) Alike (G–I), but for larger amplitude. (J) *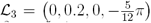*, in green; (K) 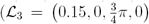, in red; (L) *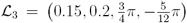*, in orange. Panel (J) and (K) also illustrate how the initial phase shifts with respect to the amplitude contribution of the negative and positive rotor depend on *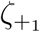*, and on *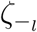* and *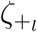*, respectively. They are given by 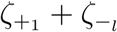 for the negative rotor and by *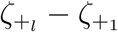* for the positive rotor.

**Figure S11:**
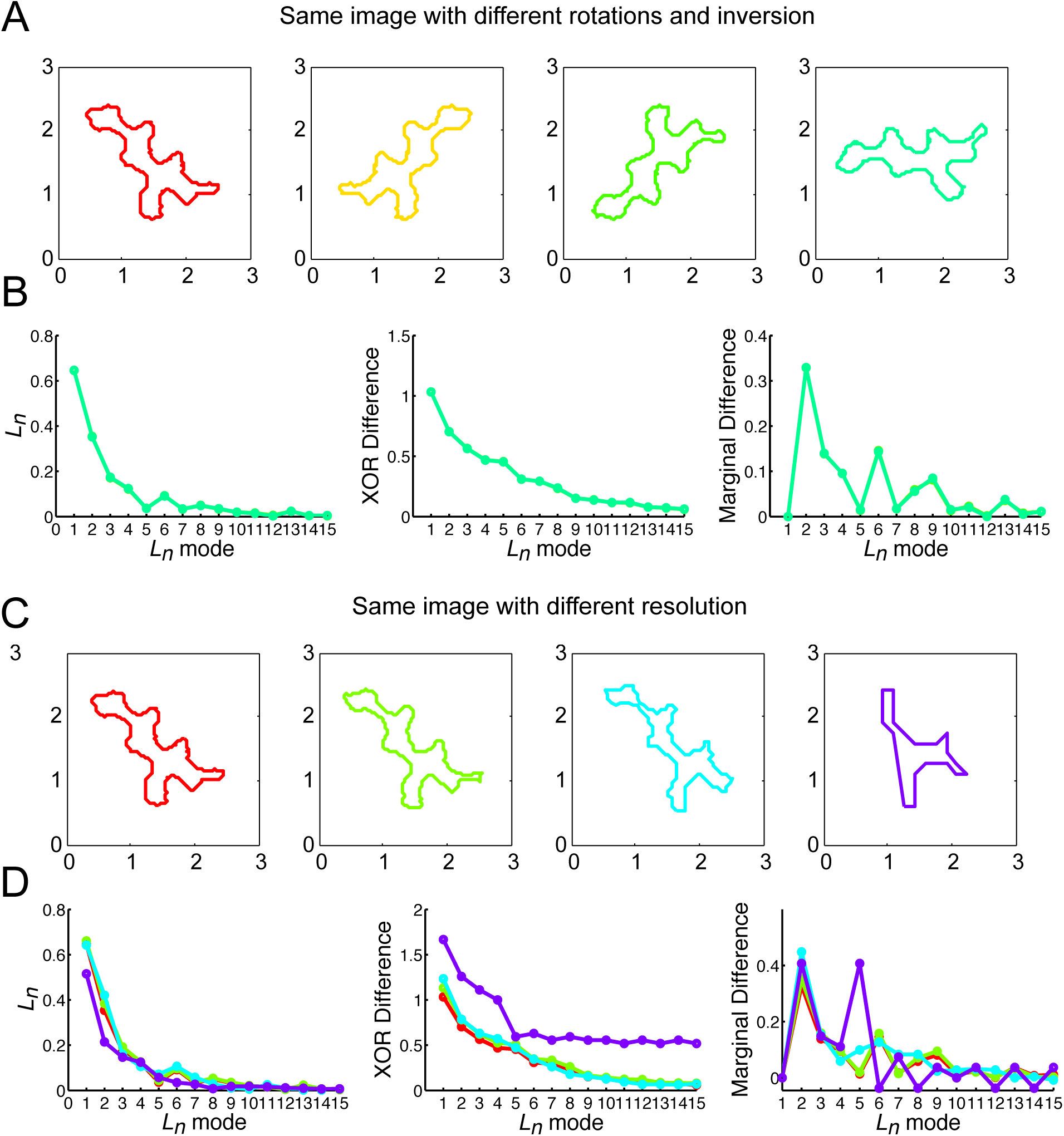
LOCO-EFA is invariant to image rotation or mirroring, nor sensitive to image resolution. (A) The cell outline of an experimentally observed cell was mirrored along the y-axis and/or rotated over different angles, after which LOCO-EFA was applied to each image separately. (B) The *L*_*n*_ values and other derived metrics were invariant to those transformations. (C) The resolution of the original image was reduced, such that the number of contour points decreased from 1104 to 253, 108 and 27 (from left to right, respectively). (D) The *L*_*n*_ numbers and the associated metrics only deviated from the high-resolution values when the resolution was very low and the cell outline was clearly deviating from original cell outline. This in contrast to the skeletonisation method, for which even marginal resolution reductions can cause large deviations in the outcome.

## Supporting Table

**Table S1:** Three types of specified cell shapes (SCS) as used in Figure S7 and Figure S8.

|  | SCS1 | SCS2 | SCS3 |
| --- | --- | --- | --- |
| Target Area | 858 | 1167 | 1197 |
| Pointedness | 6912 | 5328 | 5207 |
| Number of lobes | 3–10 | 3–10 | 3–10 |
| Roundness | 382 | 518 | 434 |
| Elongation | 4 | 28 | 5927 |

## Supporting Movies

Movie S1: **Approximation of a closed contour using Elliptic Fourier Analysis**. A given two-dimensional shape can be approximated using EFA by summing *n* elliptic harmonics as follows: each *n*th elliptic harmonic traces *n* clockwise or counter-clockwise revolutions while moving around the previous elliptic harmonic.

Movie S2: **Direction of rotation opposite to the first harmonic ellipse**. EFA mode 3 generating a shape with 4 features, its rotation direction opposite to the rotation direction of the first harmonic.

Movie S3: **Direction of rotation same as the first elliptic harmonic**. EFA mode 3 generating a shape with 2 features, its rotation direction the same as the rotation direction of the first harmonic.

Movie S4: **Exception of the rule regarding rotation direction**. If the eccentricity of an elliptical harmonic is very high, the number of generated lobes does not follow the rule-of-thumb proposed by Diaz et al. (1990). Here, EFA mode 3 generates a shape with 4 features, although its rotation direction is the same as the rotation direction of the first harmonic.

Movie S5: **Example of an *in silico* simulation of a population of cells with more complex specified shapes**. CPM simulation of multi-lobed cells take up pavement-like cell shapes when they are allowed to interact with their neighbours within a confluent tissue.

## Supporting Code

**loco-efa example.c** Code written in C which applies the full procedure to calculate the *ℒ*_*l*_ and *L*_*l*_ values to any file containing contour coordinates. For each computational step it is indicated where the relevant mathematical details can be found in the Supplementary Materials and Methods. We have intentionally kept the code as bare as possible, without for example any graphical interface, to allow it to be trivially compiled and run on any platform. Details regarding compilation and execution can be found in the header of the file.

**cell outline.csv** Contour data of the cell presented in Figure 3S, Figure S3A and Figure S11. This file, or any other file containing contour data, can be used as an input for the program.

